# Introduction of probiotic bacterial consortia promotes plant growth via impacts on the resident rhizosphere microbiome

**DOI:** 10.1101/2021.06.21.449210

**Authors:** Jie Hu, Tianjie Yang, Ville-Petri Friman, George A. Kowalchuk, Yann Hautier, Mei Li, Zhong Wei, Yangchun Xu, Qirong Shen, Alexandre Jousset

**Affiliations:** Jiangsu Provincial Key Lab for Organic Solid Waste Utilization, Key Lab of Plant immunity, National Engineering Research Center for Organic-based Fertilizers, Jiangsu Collaborative Innovation Center for Solid Organic Waste Resource Utilization, Nanjing Agricultural University, Weigang 1, Nanjing, 210095, PR China; Utrecht University, Institute for Environmental Biology, Ecology & Biodiversity, Padualaan 8, 3584CH Utrecht, the Netherlands; University of York, Department of Biology, Wentworth Way, York, YO10 5DD, United Kingdom

**Keywords:** Rhizosphere microbiome, biodiversity, probiotic consortia, multifunctionality, *Pseudomonas*, plant growth-promotion

## Abstract

Plant growth depends on a range of functions provided by their associated rhizosphere microbiome, including nutrient mineralization, hormone co-regulation and pathogen suppression. Improving the ability of plant associated microbiome to deliver these functions is thus important for developing robust and sustainable crop production. However, it is yet unclear how beneficial effects of probiotic microbial inoculants can be optimised and how they are mediated. Here, we sought to enhance the tomato plant growth by targeted introduction of probiotic bacterial consortia consisting of up to eight plant-associated *Pseudomonas* strains. We found that the effect of probiotic consortium inoculation was richness-dependent: consortia that contained more *Pseudomonas* strains, reached higher densities in the tomato rhizosphere and had clearer beneficial effects on multiple plant growth characteristics. Crucially, these effects were best explained by changes in the resident community diversity, composition and increase in the relative abundance of initially rare taxa, instead of introduction of plant-beneficial traits into the existing community along with probiotic consortia. Together, our results suggest that beneficial effects of microbial introductions can be driven indirectly through effects on the diversity and composition of resident plant rhizosphere microbiome.

## Introduction

Microorganisms associated with plant roots provide a range of services essential for plant growth. Many species mineralise nutrients, produce growth hormones or prevent diseases [1,2]. However, while many functions have been described, it has remained to date challenging to harness the multiple functions encoded in the microbiome. This is particularly an issue in soils degraded by intensive agriculture. In such soils, the low microbial biodiversity restricts the number of plant-beneficial functions and makes their expression by microbes unstable [3]. Consequently, soil fertility may be improved by restoring microbial biodiversity, thereby re-establishing community-level delivery of the multiple plant-beneficial services needed for a robust growth [4,5].

One way to increase microbiome-associated multifunctionality is the introduction of additional microbes into the soil [6]. For a long time, microorganisms have been selected on the base of their ability to directly express functions of interests [7], such as nutrient mineralisation, nitrogen fixation or pathogen suppression [8]. However, such direct effects are notoriously unstable as the introduced species fail to establish at the density needed to function in a natural microbiome context [9]. Further, inherent trade-offs in microbial physiology will limit the expression of functions one microbial species or strain can provide to the plant [10]. Such limitations could be partly solved by using multispecies probiotic consortia to improve both the inoculant establishment and the variety of services microbes can provide to the plant [11,12] compared to single-species inoculants [13–15]. For example, a consortium of multiple species that do not show antagonism towards each other, may together occupy a broader range of ecological niches [16], allowing them to better colonise plant rhizosphere. Furthermore, a more diverse microbial consortium will likely contain a larger amount of plant-beneficial functions [11,16,17], increasing the consortium-level functional diversity and redundancy. Hence, application of microbial consortia could potentially introduce multiple plant-beneficial functions into the soil that could be expressed simultaneously by different consortium members due to ecological complementarity.

In addition to directly introducing functions to the rhizosphere, inoculated microbes could have indirect effects via alteration of the diversity, composition and functioning of the resident rhizosphere microbiome [18,19]. Previous studies have shown that microbial invasions can considerably shift microbiome diversity, composition and functioning leading to significant effects even over successive plant generations [4,5]. Such changes could be mediated via microbial competition for resources or antagonism triggered by antibiosis [20], which could change the balance between dominant and rare taxa [21]. Some inoculated species could also interact with the plant, triggering plant-mediated ‘steering’ of microbiomes via altering root exudation patterns [22], producing plant-derived antimicrobials [23] or inducing other plant defences [24]. If the effects of species introductions are diversity-dependent [25], diverse microbial inoculants could trigger relatively larger shifts in the functioning of resident microbiome [26]. While such potential indirect effects of microbial inoculants on the functioning of the resident rhizosphere microbiome have been reported previously [18,19], they are still relatively poorly understood.

In this study, we employed a biodiversity-ecosystem functioning framework [27] to assess how the diverse probiotic bacterial inoculants affect rhizosphere microbiome and their impact on plant growth. To this end, we assembled probiotic bacterial consortia consisting of one to eight different *Pseudomonas* spp. strains that all have previously been shown to have beneficial effects on plants (Table S1-3) [28]. We first characterised the *in vitro* performance of each consortium regarding bacterial traits linked to plant-beneficial functions (Table S3). The mean effect of individual plant-beneficial functions were further analyzed by a weighted multifunctionality index [29], which summarised the overall consortium ability to provide different functions simultaneously. We then inoculated all probiotic consortia in the tomato rhizosphere, and tested their effects on the plants and resident microbiome. First, we measured the consortium effects on the plant growth, nutrient assimilation and protection against the soil-borne pathogen, *Ralstonia solanacearum* [30]. Second, we used 16S rRNA amplicon sequencing to obtain a snapshot of tomato rhizosphere microbiome after the inoculation of *Pseudomonas* consortia, and assessed the probiotic inoculant colonisation. Finally, structural equation modelling was used to compare the contribution of direct and indirect effects of the inoculated consortia on plants. We hypothesized that introduction of *Pseudomonas* consortia could promote plant growth, and that this promotion could be magnified with increasing consortium richness. Mechanistically, such effects could be driven by improved *Pseudomonas* establishment and introduced novel functional traits into the microbiome directly, or indirectly via changes on the composition and functioning of the existing resident microbiome.

## Material and Methods

### Model bacterial strains

We used eight strains of *Pseudomonas* spp. as model organisms: *P. fluorescens* 1m1-96, *P. fluorescens* mvp1-4, *P. fluorescens* Phl1c2, *P. fluorescens* Q2-87; *P. kilonensis* F113, *P. protegens* Pf-5 and CHA0; *P. brassicacearum* Q8r1-96 (Table S1). These strains are well-characterised model strains and widely used in plant growth-promotion and pathogen-suppression studies [17,28,31,32]. They express different traits that are important for several microbiome functions that improve plant growth (Table S3). *Ralstonia solanacearum* QL-Rs1115 strain was used as a model soil-borne pathogen in *in vitro* and *in vivo* experiment. All strains were stored at −80 °C, and prior to all experiments, one single colony of each bacterial strain was selected and prepared by grown overnight in nutrient broth medium, washed three times in 0.85 % NaCl buffer and adjusted to a density of 10^8^ cells mL^−1^.

### Assembly of probiotic *Pseudomonas* consortia

We assembled a total of 37 distinct *Pseudomonas* consortia in 48 treatments that contained 1, 2, 4 or 8 *Pseudomonas spp.* strains (four richness levels) following a substitutive design (Table S2) [17,33,34]. In this design, each strain is equally often present in different consortia within each richness level, allowing discriminating richness effects from confounding species identity effects later in the analysis [34]. Bacterial suspensions of individual *Pseudomonas* species were mixed in equal amounts in all consortia, which were immediately used for subsequent experiments. The same substitutive experimental design was used both the *in vitro* assays and *in vivo* greenhouse experiments.

### Measurement of *in vitro* functional traits of the assembled consortia and calculation of multifunctionality index

The original data related to *in vitro* functional traits of the assembled consortia were measured in previously published studies [11,17]. In this study, all previously collected data was combined for an integrated meta-analysis (Table S4) and short description of measurement details are in supplementary materials.

### Calculation of *in vitro* multifunctionality index

All measured *in vitro Pseudomonas* traits were used to compute a multifunctionality index for each consortium using a weighted average standardized calculation method [29]. Briefly, all measured variables were first correlated with the consortium richness to detect the direction of correlations, then standardized between 0 and 1 by using a modified function of the getStdAndMeanFunctions in R [35]. To avoid an overrepresentation of functions with similar contribution to overall ecosystem functioning, we applied a cluster analysis [29] to identify closely related functions. We assigned a weight of one to each cluster and weighed each function equally within each cluster so that functions in a cluster summed to one. The weighted average multifunctionality was then calculated based on the dendrogram. For example, siderophore and auxin production were assigned to one cluster and each of them received a weight of 0.5 (Fig. S1A).

### Greenhouse experiments

We assessed the effect of consortium richness on microbiome diversity and function in two separate greenhouse experiments. In both experiments, we grew tomato plants in non-sterile agricultural soil collected from a tomato field in Qilin town of Nanjing, China, which has been used for tunnel greenhouse farming for over 15 years [36]. Surface-sterilized tomato seeds (*Lycopersicon esculentum*, cultivar “*Jiangshu*”) were germinated on water-agar plates for three days before sowing into seedling plates containing ^60^Co-sterilized seedling substrate (Huainong, Huaian Soil and Fertilizer Institute, Huai’an, China). Germinated tomato plants were transplanted to seedling trays containing non-sterile agricultural soil at the three-leaf stage (12 days after sowing). Two tomato plants per cell (600 grams of soil) were transplanted to total of 8 separate cells resulting in sixteen seedlings per seedling tray (370 mm × 272 mm × 83 mm). Each tray was treated as one biological replicate resulting to a total of 104 trays. Half of these trays (N=52) were used for plant growth experiment, and the other half (N=52) for plant protection experiment, more details were in supplementary materials.

### Calculation of weighted average plant growth index

All measured plant traits (plant aboveground dry biomass; nitrogen, potassium, phosphorus and iron concentrations in the plant tissue; and plant disease severity index) were used to calculate weighted average plant growth index [29]. To avoid overrepresentation of functions with similar contribution, we applied the cluster analysis to all measured traits similar to when previously calculating *in vitro Pseudomonas* consortium multifunctionality index. For example, aboveground plant biomass and protection against pathogen were assigned to one cluster and each of them received a weigh of 0.5 (Fig. S1B).

### DNA extraction, qPCR quantification and sequencing

#### Soil DNA extraction

Rhizosphere soil from the plant growth promotion experiment was collected by gently removing plants from the trays before shaking off excess soil and collecting the soil attached to the root system. Two randomly selected plants from different cells per seedling tray were collected and pooled together. Samples were then suspended into 30 mL of sterile H_2_O (100 rpm, 30 min at 4 °C) and centrifuged (5000 g, 30 min at 4 °C) before the soil pellets were transferred into 2 mL tubes and stored at −80 °C for subsequent experiments. We extracted DNA with the Power Soil DNA Isolation Kit (Mobio Laboratories, Carlsbad, CA, USA). Briefly, DNA from 0.3 g of soil pellets per sample was extracted following the manufacturer’s protocol. DNA fragment size was checked on 1% agarose gel, and DNA concentration and purity were determined with a NanoDrop 1000 spectrophotometer (Thermo Scientific, Waltham, MA, USA) prior to downstream analyses.

#### Quantification of total bacterial and Pseudomonas abundances

We used quantitative PCR (qPCR) to quantify the abundance of total bacteria associated with plant rhizosphere soil as well as the introduced *Pseudomonas* consortia based on 16S rRNA and *phlD* gene copies per gram of soil, respectively. More details were in supplementary materials.

#### 16S rRNA amplicon sequencing

We used the 563F (5’-AYT GGG YDT AAA GVG-3’) and 802R (5’-TAC NVG GGT ATC TAA TCC-3’) [37] primer pair to amplify the V4 hypervariable region of the bacterial 16S rRNA gene, more detailed description was in supplementary materials. Raw fastq files were demultiplexed and quality-filtered with QIIME (version 1.17) [38] according to previous established protocols [19]. After discarding unqualified reads, the operational taxonomic units (OTUs) were assigned at 97% identity level using UPARSE [39] and chimeric sequences identified and removed using UCHIME [40]. The phylogenetic affiliation of each 16S rRNA gene sequence was analyzed using the RDP Classifier [41] against the Silva 16S rRNA gene database with a confidence threshold of 70% [42]. We removed unassigned, Archaea, mitochondrial and plastid OTUs and those found in fewer than 3 times in less than 1% of the samples. PyNast and FastTree were used to estimate the phylogeny of all OTUs observed in all samples. To obtain an equivalent sequencing depth for later analysis, the sample OTU abundances were rarefied by using the lowest sample sequence depth, resulting in similar sequence depths between all samples (mean=29,068, min=28,749, max=29,465). To ensure the robustness of the OTU bioinformatic pipeline, the 16S rRNA Miseq sequence data was also analyzed using DADA2 pipeline. More details were included in supplementary materials. See Fig. S2 for a reproduction of Fig. 4A, and Fig. S3 for a reproduction of Fig. 4C with DADA2 pipeline.

#### Calculation of resident microbiome biodiversity

We characterized the biodiversity of resident bacterial communities using the phylogenetic abundance evenness (PAE), an index accounting for both evenness and phylogenetic distribution, which was calculated by [43], more details were in supplementary material and methods.

### Statistical analyses

To analyze the effect of richness and abundance of probiotic *Pseudomonas* consortia on the resident rhizosphere microbiome community composition, diversity and weighted mean of plant growth index, generalised linear models (GLMs) were used. Consortium richness was treated as a factor, *phlD* abundance in rhizosphere was log-transformed, and their effects on subsequently measured parameters were assessed with GLMs using a Gaussian distribution in two models with different factor directions following a subsequent ANOVA analysis in R.

The effect of microbial inoculation on rhizosphere microbiome community composition was determined with a Redundancy analysis (RDA) based on the taxa presence-absence OTU table, consortium richness (factor with five levels) and *phlD* gene copy numbers in the rhizosphere, using the R function vegan: rda. Coordinates were used to draw 2D graphical outputs and the effect of consortium richness and establishment success on rhizosphere microbiome community composition was tested using the R function vegan: adonis.

To explore the significance of *Pseudomonas* strain properties for the shift in resident microbiome, the effect of *in vitro* plant-beneficial functions measured at the consortium level were used to explain the variation in the resident rhizosphere microbiome composition which was presented by using canonical correlation analysis. Stepwise GLMs were used to test the effect of *in vitro* plant-beneficial functions, such as breadth of carbon catabolism, antibacterial activity, phytohormone production and nutrient availability, on resident microbiome composition.

Finally, we used structural equation modelling (SEM) to examine direct and indirect effects linking consortium inoculants with plant growth by accounting for multiple potentially correlated effect pathways [44]. The initial model was based on reported data on plant-microbe interactions and microbial ecology [45], assigning the “richness” as exogenous variables and “*in vitro* multifunctionality index”, “microbiome community composition”, and “weighted average of plant growth index” as endogenous variables, respectively. The adequacy of the models was determined via χ^2^-tests and Akaike information criterion (AIC) [45]. Structural equation modelling was performed using the R package piecewiseSEM.

## Results

### Effect of richness and strain identities on probiotic consortium plant-beneficial functions

Each of the studied *Pseudomonas* strain excelled at expressing a specific subset of plant-beneficial functions, including ability to consume different carbon resources, production of auxin, gibberellin, siderophores or antibacterial activity (Fig. 1A). For example, *Pseudomonas protegens* Pf-5 produced the highest amount of siderophores but the lowest levels of auxin (Fig. 1A). In contrast, *Pseudomonas fluorescens* 1M1-96 produced the highest levels of auxin but had a very weak antibacterial activity (Fig. 1A). These differences suggest that *Pseudomonas* strains were specialized in different plant-beneficial functions and could potentially show complementary effects when combined in multi-strain consortia. To test this, we used weighted means of all the individual plant-beneficial functions and combined them into predicted consortium multifunctionality index. It was found that consortium richness correlated positively with the consortium multifunctionality index (Fig. 1B, F_1, 46_ = 60.02, P < 0.0001), and increasing the consortium richness promoted the expression of all the individual plant-beneficial functions when measured directly *in vitro* (Table S5). Some of the individual functions were further influenced by the presence of certain strains. For instance, consortia containing the *Pseudomonas protegens* CHA0 and Pf-5 strains showed the highest levels of antibacterial activity (Table S5). However, despite certain strong identity effects, consortium richness remained a major significant predictor even after accounting for the inclusion of each strain.

**Figure 1.**
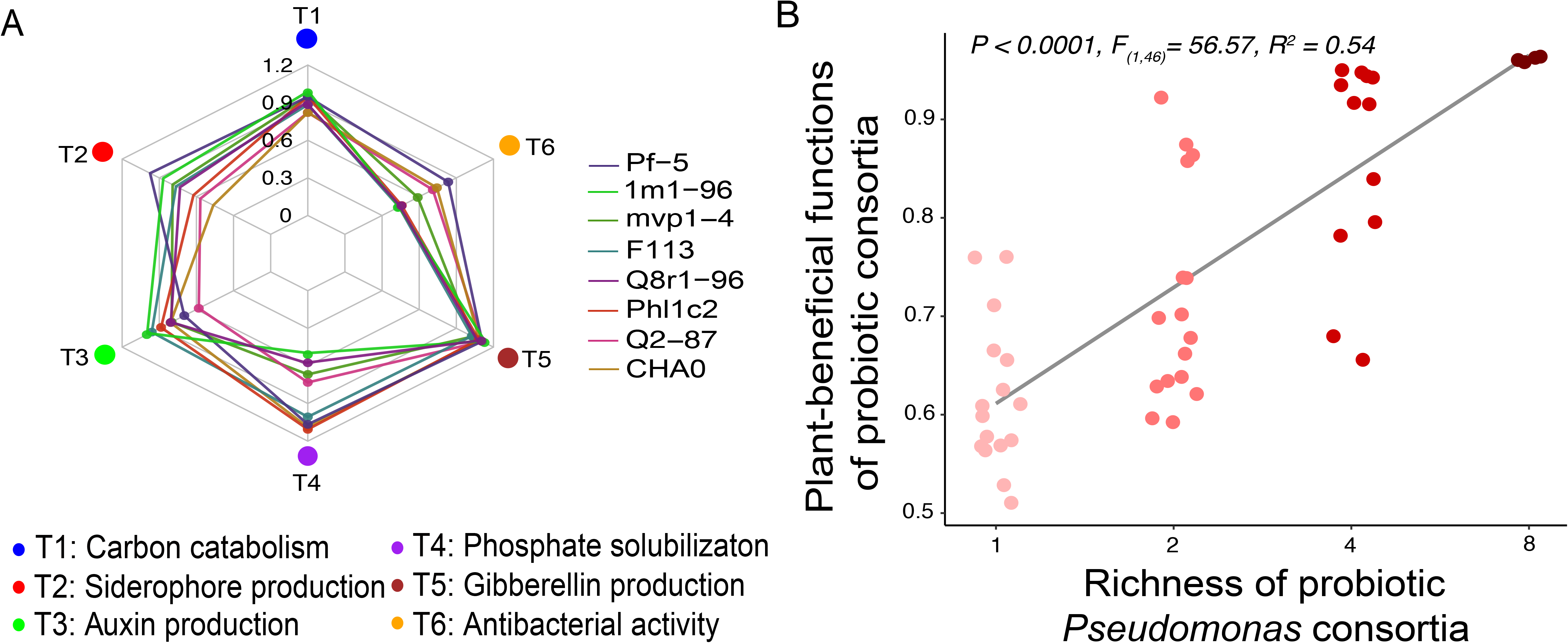
*in vitro* plant-beneficial functions provided by the probiotic *Pseudomonas* strains and consortia. A: Radar chart showing the mean ability of individual *Pseudomonas* strains of to express six plant-beneficial functions measured *in vitro*. B: Positive correlation between the richness of probiotic *Pseudomonas* consortia and mean of all plant-beneficial functions measured *in vitro* at the consortium level (consortia assembled based on substitutive design presented in Table S2).

### Effect of consortium richness and strain identities on plant growth characteristics

Similar to *in vitro* measurement of plant-beneficial functions, each *Pseudomonas* strain had unique impacts on plant growth on their own. For example, *Pseudomonas kilonensis* F113 inoculation led to a high aboveground plant biomass, *Pseudomonas protegens* CHA0 inoculation led to a high concentration of potassium and phosphorous in plant tissues, while *Pseudomonas brassicacearum* Q8r1-96 inoculation led to the highest protection against pathogen (Fig. 2A). Single *Pseudomonas* strain inoculations affected only a limited number of plant growth characteristics (Fig. 2A-B), increasing consortium richness led to an improvement of multiple plant growth characteristics (Fig. 2B). All individual plant growth characteristics were positively correlated with consortium richness except for plant tissue nitrogen concentration (Fig. 2B). As a result, also the weighted average plant growth index correlated positively with the probiotic consortium richness (Fig. 2B). In contrast to the *in vitro* plant-beneficial functions expressed by consortia, the *Pseudomonas* strains had no significant identity effects on the plant growth (Fig. 2; Table S5).

**Figure 2.**
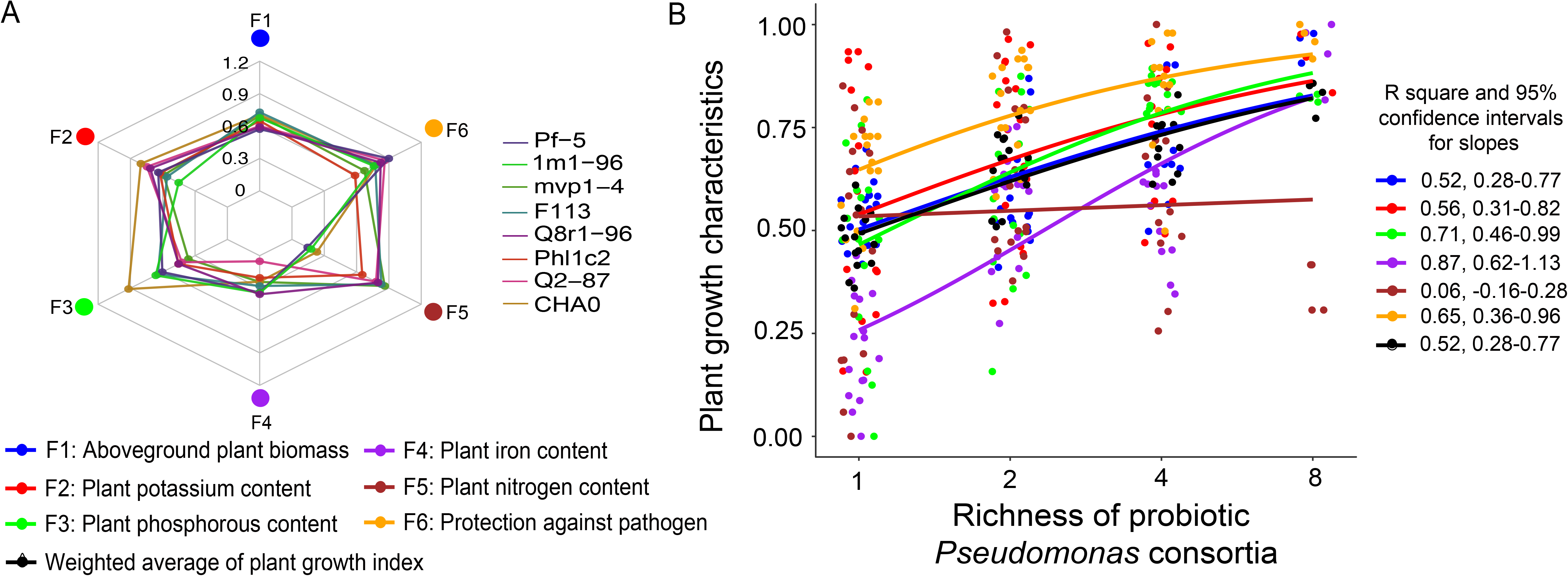
Plant growth benefits provided by the introduced *Pseudomonas* strains and consortia measured *in vivo*. A: Radar chart showing the mean effect of individual *Pseudomonas* strains on six plant growth characteristics. B: Correlation between the richness of probiotic *Pseudomonas* consortia and the six individual plant growth characteristics (colored lines), and the weighted average of plant growth index (black line) by probiotic consortia (consortia assembled based on substitutive design presented in Table S2; each point shows the mean effects of three replicate plants (N=3) for each plant growth characteristic when exposed to different consortia). The colors in the key of panel A correspond to the same growth characteristics and colors presented in panel B.

### Effect of richness and strain identities on consortium establishment in the rhizosphere

The background *phlD* gene abundances were below 0.001% (relative to 16S rRNA gene abundances) in the non-inoculated control treatments (Fig. 3A), confirming that *Pseudomonas* strains containing this gene were rare in the non-sterile agricultural soil used in the pot experiments. In contrast, *phlD* gene abundances ranged between 0.01% and 0.25% of the total 16S rRNA gene abundances in samples with inoculated *Pseudomonas* strains. Specifically, we found that *phlD* gene abundances increased along with consortium richness gradient (Fig. 3A). Only mvp1-4 strain had a marginally significant positive identity effect on the relative *phlD* gene abundances (P = 0.04, Table S6).

**Figure 3.**
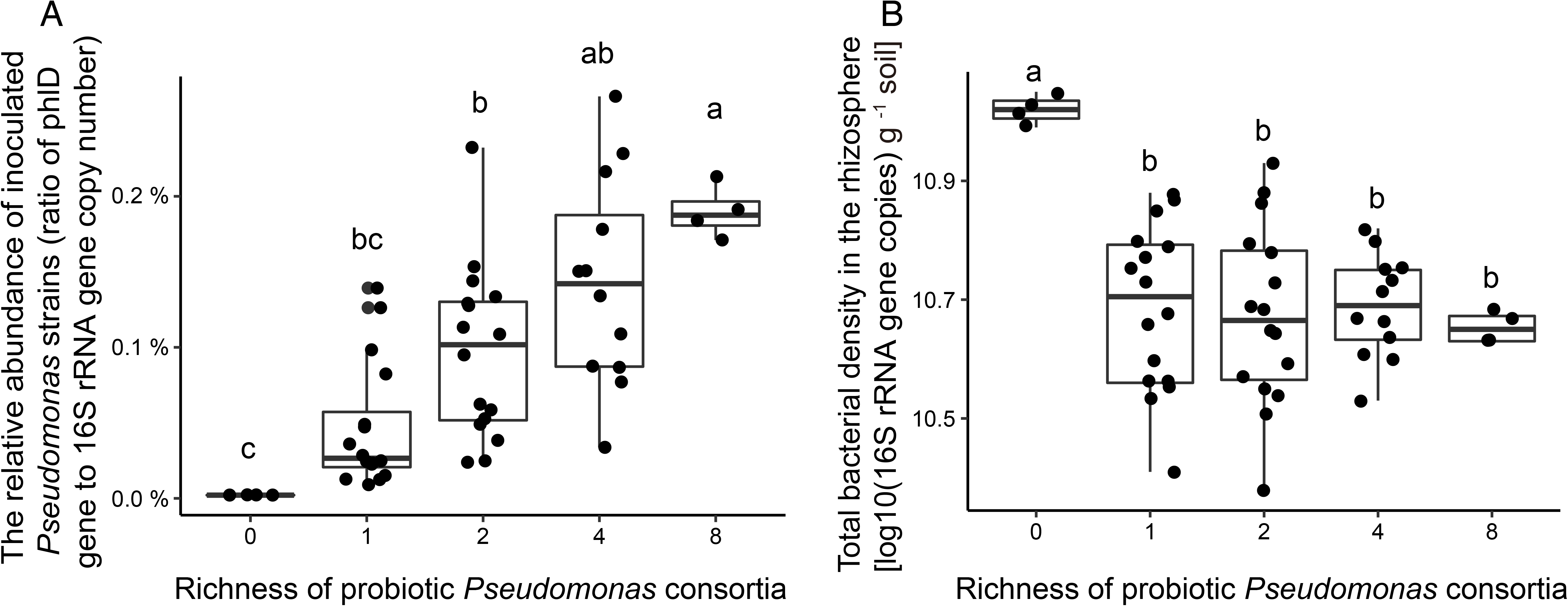
Relationship between probiotic consortia richness and the relative *Pseudomonas* (A) and total bacterial abundances (B) in the tomato rhizosphere microbiome. A: The relative mean abundances of *Pseudomonas* strains at different consortium richness levels (measured as the *Pseudomonas*-specific *phlD* gene copy numbers relative to total 16S rRNA gene copy numbers). B: The total bacterial densities at different *Pseudomonas* consortium richness levels (quantified as the total 16S rRNA gene copy numbers per gram of rhizosphere soil; presented on a log10-scale). In both panels, each point corresponds to a different consortium. Duncan’s New Multiple Range Test was used to compare differences between richness levels, and significances are denoted by different letters above the boxplots.

### Effect of consortium richness on the abundance, composition and diversity of resident rhizosphere microbiome

We found that irrespective of inoculant richness, all probiotic consortia had similar negative effects on the abundance of resident rhizosphere bacteria compared to non-inoculated control treatment (Fig. 3B). All inoculated consortia changed the composition of the resident rhizosphere microbiome. However, this impact was magnified with increasing consortium richness (P = 0.0280, Table 1; Fig. 4A). Interestingly, a clear increase in resident microbiome diversity was observed along with consortium richness (Table 1 and Table S7), even though no differences between single- and two-strain consortium, or four- and eight-strain consortium inoculants were found (Fig. 4B). Moreover, the effects of consortium richness on microbiome diversity remained significant even after accounting for *Pseudomonas* strain identity effects (Table S7). One explanation for the increase in the resident community diversity is that *Pseudomonas* inoculants potentially revived rare dormant species. In support for this, we found that rare taxa at phylum, family and OTU levels were more likely to increase in abundance along with the richness of the inoculated consortia (χ^2^ _(1,8607)_ = 22.45, P < 0.0001, Fig. 4C, Fig. S4A-C). In contrast, the most abundant taxa were more likely to decrease in response to *Pseudomonas* inoculations (χ^2^ _(1,8607)_ = 6.51, P < 0.0001, Fig. 4C, Fig. S4D). At a coarser taxonomic level, we observed that resident microbiome responses to *Pseudomonas* inoculants were conserved with some phyla. For instance, OTUs belonging to the phyla Candidate division WS6, Thermotogae, Spirochaetae, Gracilibacteria and Parcubacteria consistently increased along with the *Pseudomonas* consortium richness (Fig. S4A, Table S8). Finally, we conducted co-variance network analysis to assess whether the introduced *Pseudomonas* strains were embedded in the resident microbiome. Despite the limited taxonomic resolution of amplicon sequencing, we could identify two *Pseudomonas sp.* OTUs (shown as green circles in modules 1 and 2, Fig. 4D) putatively matching with two introduced *Pseudomonas* strains (CHA0 and mvp1-4 strains). These OTUs covaried positively with a range of other OTUs present in the resident rhizosphere microbiome, including Parcubacteria, Proteobacteria, Actinobacteria, Bacteroidetes and Saccharibacteria phyla (Fig. 4D; Table S9).

**Figure 4.**
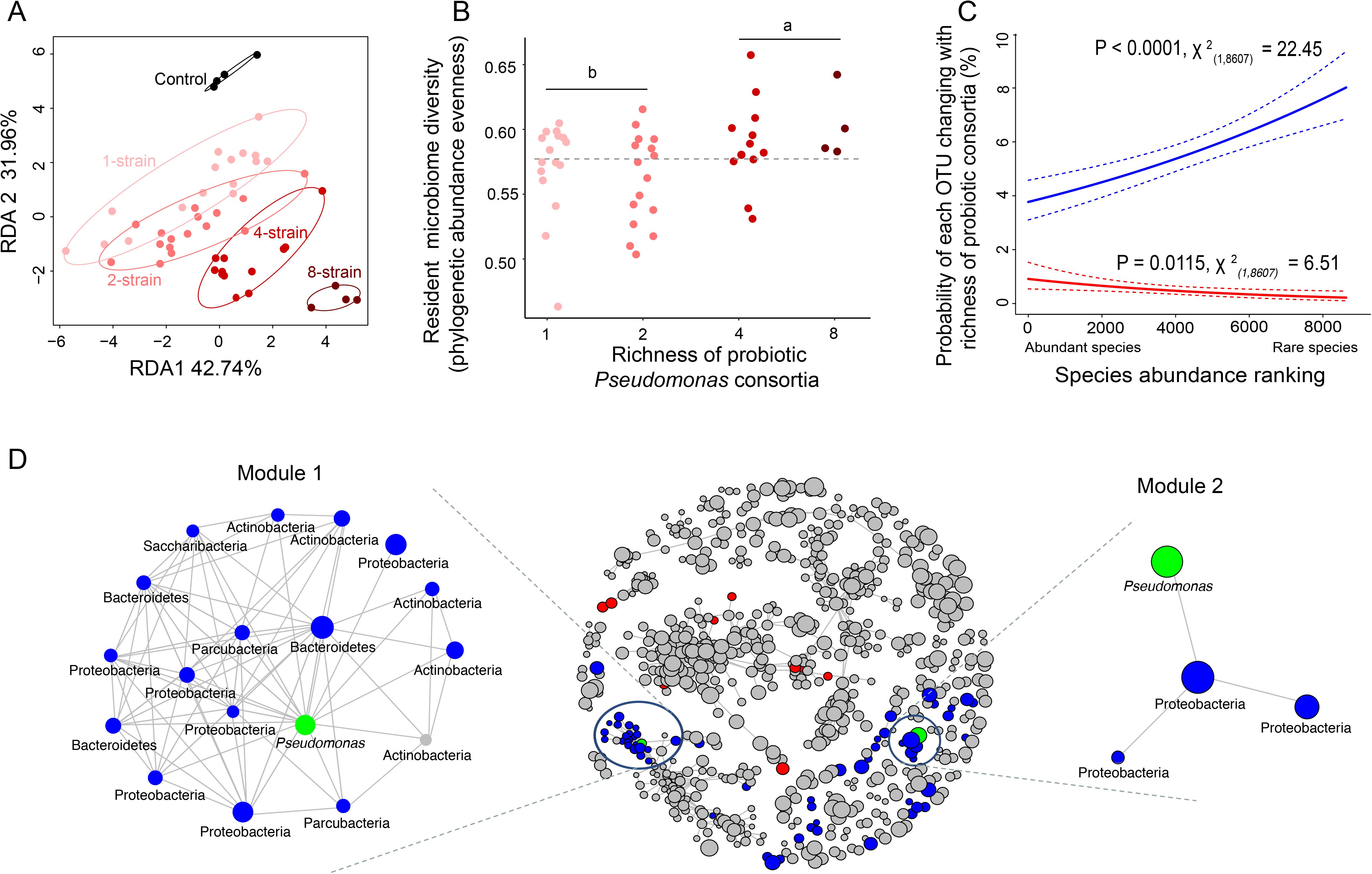
Effects of probiotic *Pseudomonas* consortia on the composition and diversity of the resident rhizosphere microbiome. A: RDA analysis of resident microbiome composition showing the impact of increasing consortium richness on the rhizosphere microbiome composition. Ellipses enclosing points at each richness level show 95 % confidence intervals. B: Effect of probiotic consortium richness on the rhizosphere microbiome diversity in terms of phylogenetic abundance evenness. The grey dashed line shows the mean diversity in the control treatment (non-sterile agricultural field soil without introduced consortia). Tukey HSD analysis was performed to compare low-diversity (one- and two-strain) and high-diversity (four- and eight-strain) consortia. In both panels A and B, each point corresponds to different consortium (based on substitutive design presented in Table S2). C: response of abundant and rare rhizosphere microbiome taxa to increasing richness of probiotic consortia. The blue line shows the binomial regression on the probability of a given OTU (Y-axis), which was initially abundant or rare (X-axis, OTUs were ranked based on their relative abundance in the control treatments), to increase in its relative abundance along with consortium richness. The red line shows the binomial regression on the probability of a given OTU (Y-axis), which was initially abundant or rare (X-axis), to decrease in its relative abundance along with consortium richness. Dashed lines show 95% confidence interval of each binomial regression. D: Co-occurrence network of the resident microbiome (in the middle), with each node representing a bacterial OTU where the node size is proportional to relative OTU abundance. Blue and red circles represent OTUs that were significantly positively and negatively correlated with consortium richness, respectively. Green circles indicate OTUs (OTU 8478 in module 1 and OTU4923 in module 2) putatively belonging to the inoculated CHA0 and mvp1-4 strains. Links between nodes show statistically significant positive Spearman correlations (P < 0.05) with corelation coefficient > 0.8 (no significant negative correlations found). Left and right insets show detailed overview of modules 1 and 2, highlighting the associations between resident taxa and putative probiotic *Pseudomonas* OTUs (OTUs are shown at lowest taxonomic level that could be assigned by the bioinformatic analysis, ranging from phylum to genus level).

### Disentangling direct vs. indirect effects of consortium richness on plant growth

We used Structural Equation Modelling (SEM) to disentangle whether the effects of probiotic consortia on the plant growth were directly driven by the plant-beneficial functions introduced by consortia, or indirectly via shifts in the resident rhizosphere microbiome. We found that plant-beneficial functions of consortia measured *in vitro* predicted poorly the individual plant growth characteristics (Fig. S5A-C). Only notable exception was a positive relationship between the consortium phosphate solubilization capacity and the plant shoot phosphorus content (Fig. S5D). Similarly, the multifunctionality index of probiotic consortia did not explain the positive effects of inoculants on the plant growth (left side, Fig. 5). Instead, *Pseudomonas* consortium richness had strong effects on the composition of resident plant microbiome, which in turn was positively correlated with weighted average plant growth index (Fig. 5). Interestingly, consortium diversity had also a direct positive effect on weighted average plant growth index (residuals; Fig. 5).

**Figure 5.**
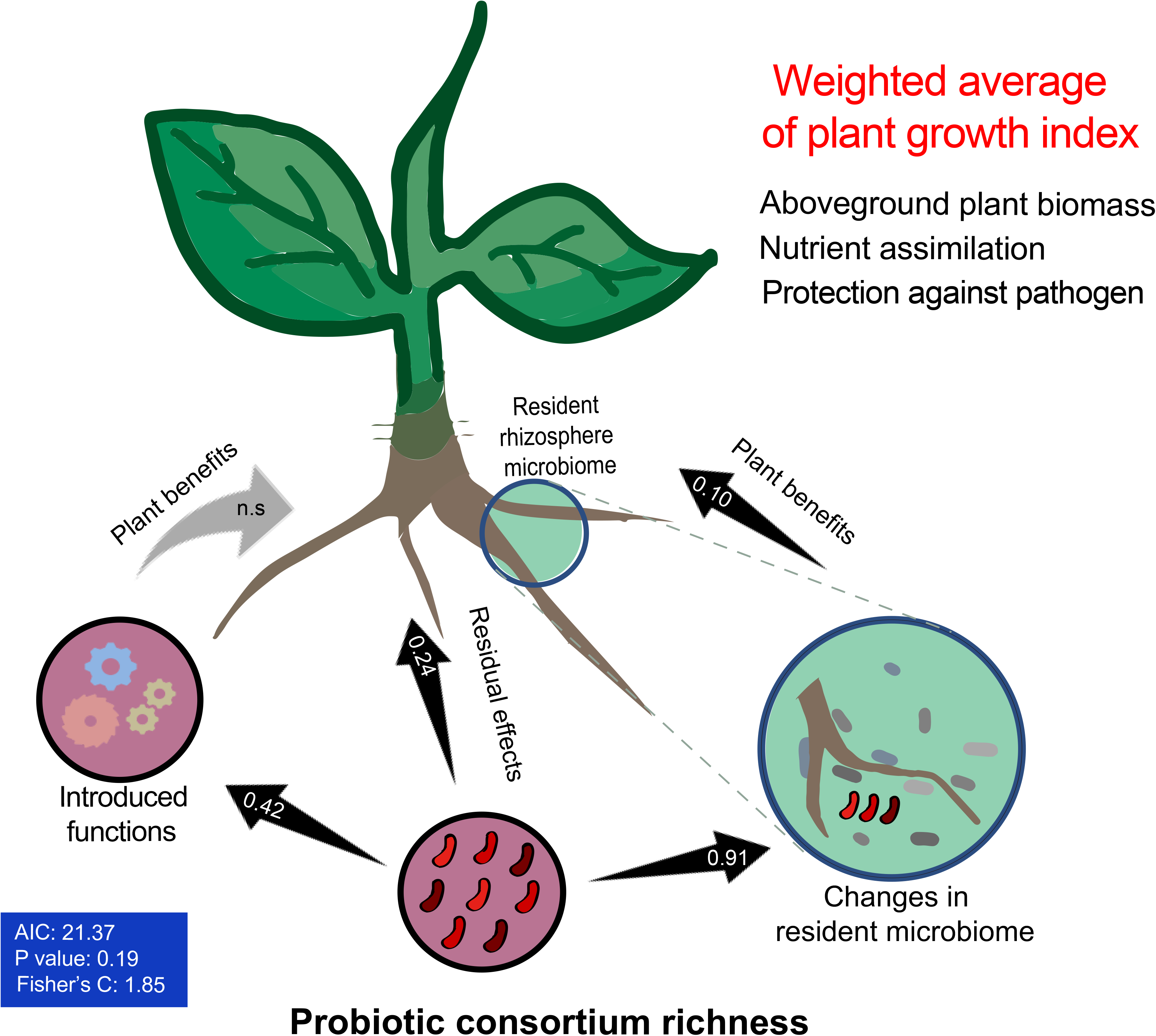
Structural equation model (SEM) comparing the direct and indirect effects of consortium richness on plant growth (weighted average of plant growth index). Black arrows indicate significant relationships between the tested variables, while grey arrows indicate non-significant relationships retained in the model. ‘Introduced functions’ refer to consortium multifunctionality index, while ‘Changes in resident microbiome’ refers to shift in the resident microbiome composition after inoculation (the first RDA axis of multivariate community composition analysis). The numbers inside the arrows indicate standardized correlation coefficients (relative effect sizes of non-significant correlations are not shown). The parameters inside the blue rectangle on the left show the robustness of the SEM model.

To gain additional insight into underlying mechanisms, we used canonical correlation analysis (CCA) to describe the resident microbiome composition and assessed if *Pseudomonas* consortium traits measured *in vitro* explained the resident microbiome composition. We found that the variation on CCA1 was best explained by the *in vitro* production of auxin and gibberellin phytohormones by the introduced consortia (Table S10). In contrast, variation on the CCA2 was best explained by the niche breadth of the probiotic consortia, *i.e.*, their ability to metabolise an array of carbon sources typically found in the rhizosphere. As access to resources is crucial for inoculant establishment, this data suggests that probiotic consortia could have shaped the resident microbiome through resource competition. In contrast, the antibacterial activity of *Pseudomonas* consortia was not retained in the final model (Table S10).

## Discussion

Here we used a biodiversity-ecosystem functioning framework to directly assess how the diversity of probiotic bacterial inoculants affect microbiome structure and plant growth. Specifically, we explored if the potential benefits were driven directly by introduced consortia via introduction of essential functions to the microbiome, or indirectly via changes in the resident bacterial community. It was found that increasing probiotic consortium richness increased inoculant colonisation success, and was associated with improved plant growth, nutrient assimilation and protection from pathogen infection. Crucially, inoculants caused shifts in the resident microbiome and these effects were magnified with increasing probiotic consortium richness, leading to an increase in the abundance of rare taxa and overall microbiome biodiversity. While some significant *Pseudomonas* strain identity effects were found, the improvement of plant growth was poorly explained by plant beneficial functions provided by different consortia members. Instead, positive effects on the plant growth were best explained by consortium-mediated shifts in the resident microbiome, which were associated with phytohormone production and resource competition by the probiotic consortia. Together these findings suggest that probiotic bacteria can be used to steer the existing resident rhizosphere microbiome in order to improve plant growth.

Multi-strain consortia could perform better together due to ecological complementarity between plant beneficial functions they can provide at consortium-level, or due to other emergent diversity effects arising in complex microbe-microbe-plant communities [20]. While the probiotic consortium richness correlated positively with its multifunctionality and several plant growth characteristics, but these consortium multifunctionality poorly predicted the plant growth. Instead, positive diversity effects via shifts in the resident microbiome and the probiotic consortium richness itself are better predictors. Overall, probiotic consortium richness effects could be explained by improved establishment success in terms of relative *Pseudomonas* abundances in the rhizosphere. While certain *Pseudomonas* strains had strong identity effects, the effect of consortium richness remained significant even after their removal. Moreover, the densities of four and eight strains consortia were up to ten times higher than the best-performing single-strain inoculants, which indicates that richness effects were driven by emergent consortium level effects. Such unexpectedly strong community performance is called transgressive overyielding [46], and could be indicative of synergistic interactions between inoculated *Pseudomonas* strains, resident microbiome or with the plant. The fact that inoculation of probiotic bacteria resulted in reduction in the abundance of resident bacteria suggests that *Pseudomonas* bacteria likely promoted competition with the resident community. In support for this, the consortium resource niche breadth increased along with the richness gradient, which might have allowed strains to sequester nutrients more efficiently in the rhizosphere [32]. However, the reduction in the resident bacteria abundances was the same for all consortia, and hence, resource competition alone is unlikely to explain the observed relationship with the *Pseudomonas* consortium richness.

Interestingly, probiotic consortium richness and establishment success were positively associated with relatively larger and non-random changes in the diversity and composition of resident rhizosphere microbiome, with 4- and 8-strain consortia showing the clearest effects. Specifically, diverse probiotic consortia were associated with more even resident bacterial communities and clearer increase in the abundance of rare bacterial taxa. One potential explanation for this could be inhibition of dominant organisms via production of secondary metabolites by the introduced consortia [47–49]. While more work is required to unravel exact mechanisms, increase in the relative abundance of rare taxa was associated with decrease in the abundance of dominant taxa (Fig. 4C), indicative of highly asymmetric responses by different resident microbiome taxa. For example, 4- and 8-strain probiotic consortia had proportionally large, positive effect on *Parcubacteria* [50], a phylum known to lack several genes responsible for the biosynthesis of essential metabolites making it likely metabolically dependent of other organisms [50]. As these organisms are still largely unknown, their potential functional role remains to be clarified in future. However, our results together suggest that the introduced *Pseudomonas* bacteria may have functioned as a keystone group reactivating the pool of rare or dormant species and their associated functions in the soil [51]. The rare rhizosphere microbiome is thus likely an important and underestimated source for beneficial bacteria [51].

In addition to resource catabolism, *Pseudomonas* phytohormone production also had significant effect, which suggest potential complex microbiome-plant feedback mediated by hormonal signalling [22,52]. For example, it has been recently shown that the presence of certain *Variovax* bacterial strains in synthetic rhizosphere communities can restore the root growth in *Arabidopsis thaliana* via effects on bacterially produced ethylene and auxin in the rhizosphere [52]. Moreover, the presence of certain bacterial taxa can modulate the expression of scopoletin antibacterial compounds by the plant via affecting root-specific transcription factor MYB72, further shaping the assembly of rhizosphere microbiome [23]. Finally, several bacteria can play an important indirect role for pathogen suppression by boosting the plant immunity instead of having direct antagonistic effect on the pathogen, including *R. solanacearum* [53]. We found similar indirect evidence in our structured equation models: consortia-mediated effects were channelled into plan growth via effects on the resident microbiome, instead of introduction of plant-beneficial functions. Our findings are thus in line with previous findings, and suggest that probiotic inoculants could be designed to reactivate the functioning of existing resident microbiomes instead of bioaugmentation of communities with additional species with desired functional traits [19]. However, as our amplicon sequencing data cannot reliably distinguish the relative abundances of different inoculated *Pseudomonas* strains, or how changes in the resident community diversity and composition were linked to expression of plant growth-promoting genes, -omics approaches and directly tailored mechanistic experiments are needed in the future. Furthermore, metabolomic and plant transcriptomics studies would be important for inferring the chemical signalling between the plant and rhizosphere microbiome [52].

## Conclusion

We conclude that plant growth can be improved using species-rich inoculants that have indirect beneficial effects for the plant through compositional changes in the resident rhizosphere microbiome. Especially, we observed positive response by rare taxa, highlighting the importance of rare biosphere for plant-microbe interactions [51]. This calls rethinking of the traditional microbial inoculant design [8]. Instead of attempting to introduce “plant-growth promoting” traits into the resident communities through microbial inoculants [54], we propose an alternative strategy where inoculants are designed to steer, boost and reactivate the resident plant growth-promoting microbes already in the rhizosphere. While more work is required to identify key bacterial taxa, their functional roles and chemical interplay with plants and other microbes [52,55], it is important to move beyond only single strain inoculants as microbe-mediated plant beneficial effects in agricultural environments are determined by complex community level interactions. This could be achieved by bringing together community ecology and -omics techniques to harness microbe-plant interactions for sustainable agriculture.

## Data Availability

The Miseq 16S rRNA sequencing data was deposited into the NCBI Sequence Read Archive (SRA) database under the accession number SRP132352. Relevant data to this manuscript and script employed in the computational analyses and plotting figures are available at https://github.com/HuJamie/JieHu2021_Microbiome.

## Statement of authorship

A.J., J.H., V.P.F., and Z.W. developed the ideas and designed the experimental plans. J.H., M.L., T.Y., Z.W. performed the experiments. J.H. and Z.W collected the data. J.H., A.J., Y.H. and Z.W. analyzed the data. A.J., J.H., Z.W., and V.P.F. wrote the manuscript. Q.S., G.K., and Y.X. gave constructive comments on the manuscript writing. The authors declare that they have no competing interests.

## Acknowledgements

We thank Shaohua Gu and Xiaofang Wang for assisting part of the greenhouse experiment, Shanghai BIOZERON Biotechnology Co., Ltd for technical support in Miseq sequencing, and Peter Veenhuizen from Utrecht University for technical support of analyzing Miseq sequencing data.

## References

1. Berendsen RL, Pieterse CMJ, Bakker PAHM. 2012 The rhizosphere microbiome and plant health. Trends in Plant Science 17, 478–486. (doi:10.1016/j.tplants.2012.04.001)

2. Vandenkoornhuyse P, Quaiser A, Duhamel M, Le Van A, Dufresne A. 2015 The importance of the microbiome of the plant holobiont. New Phytologist 206, 1196–1206. (doi:10.1111/nph.13312)

3. Liiri M, Häsä M, Haimi J, Setälä H. 2012 History of land-use intensity can modify the relationship between functional complexity of the soil fauna and soil ecosystem services – A microcosm study. Applied Soil Ecology 55, 53–61. (doi:10.1016/j.apsoil.2011.12.009)

4. Delgado-Baquerizo M, Maestre FT, Reich PB, Jeffries TC, Gaitan JJ, Encinar D, Berdugo M, Campbell CD, Singh BK. 2016 Microbial diversity drives multifunctionality in terrestrial ecosystems. Nature Communications 7, 10541. (doi:10.1038/ncomms10541)

5. Mueller UG, Sachs JL. 2015 Engineering Microbiomes to Improve Plant and Animal Health. Trends in Microbiology 23, 606–617. (doi:10.1016/j.tim.2015.07.009)

6. Zuluaga MYA et al. 2021 Inoculation with plant growth-promoting bacteria alters the rhizosphere functioning of tomato plants. Applied Soil Ecology 158, 103784. (doi:10.1016/j.apsoil.2020.103784)

7. Compant S, Samad A, Faist H, Sessitsch A. 2019 A review on the plant microbiome: Ecology, functions, and emerging trends in microbial application. Journal of Advanced Research 19, 29–37. (doi:10.1016/j.jare.2019.03.004)

8. Lugtenberg B, Kamilova F. 2009 Plant-growth-promoting rhizobacteria. Annual Review of Microbiology 63, 541–556. (doi:10.1146/annurev.micro.62.081307.162918)

9. Diallo S, Crépin A, Barbey C, Orange N, Burini J-F, Latour X. 2011 Mechanisms and recent advances in biological control mediated through the potato rhizosphere: Biological control mediated through the potato rhizosphere. FEMS Microbiology Letters 75, 351–364. (doi:10.1111/j.1574-6941.2010.01023.x)

10. Winter C, Bouvier T, Weinbauer MG, Thingstad TF. 2010 Trade-Offs between Competition and Defense Specialists among Unicellular Planktonic Organisms: the “Killing the Winner” Hypothesis Revisited. Microbiology and Molecular Biology Reviews◻: MMBR 74, 42–57. (doi:10.1128/MMBR.00034-09)

11. Hu J, Wei Z, Weidner S. 2017 Probiotic Pseudomonas communities enhance plant growth and nutrient assimilation via diversity-mediated ecosystem functioning. Soil Biology and Biochemistry 113, 122–129. (doi:10.1016/j.soilbio.2017.05.029)

12. Assainar SK, Abbott LK, Mickan BS, Whiteley AS, Siddique KHM, Solaiman ZM. 2018 Response of Wheat to a Multiple Species Microbial Inoculant Compared to Fertilizer Application. Frontiers in Plant Science 9, 1601. (doi:10.3389/fpls.2018.01601)

13. Sikes BA, Powell JR, Rillig MC. 2010 Deciphering the relative contributions of multiple functions within plant–microbe symbioses. Ecology 91, 1591–1597. (doi:10.1890/09-1858.1)

14. Aerts R, Honnay O. 2011 Forest restoration, biodiversity and ecosystem functioning. BMC Ecology 11, 29. (doi:10.1186/1472-6785-11-29)

15. Agaras BC et al. 2015 Quantification of the potential biocontrol and direct plant growth promotion abilities based on multiple biological traits distinguish different groups of Pseudomonas spp. isolates. Biological Control 90, 173–186. (doi:10.1016/j.biocontrol.2015.07.003)

16. Wei Z, Yang T, Friman V-P, Xu Y, Shen Q, Jousset A. 2015 Trophic network architecture of root-associated bacterial communities determines pathogen invasion and plant health. Nature Communications 6, 8413. (doi:10.1038/ncomms9413)

17. Hu J et al. 2016 Probiotic diversity enhances rhizosphere microbiome function and plant disease suppression. mBio 7, e01790–16. (doi:10.1128/mBio.01790-16)

18. Castro-Sowinski S, Herschkovitz Y, Okon Y, Jurkevitch E. 2007 Effects of inoculation with plant growth-promoting rhizobacteria on resident rhizosphere microorganisms. FEMS Microbiology Letters 276, 1–11. (doi:10.1111/j.1574-6968.2007.00878.x)

19. Xiong W, Guo S, Jousset A, Zhao Q, Wu H, Li R, Kowalchuk GA, Shen Q. 2017 Bio-fertilizer application induces soil suppressiveness against Fusarium wilt disease by reshaping the soil microbiome. Soil Biology and Biochemistry 114, 238–247. (doi:10.1016/j.soilbio.2017.07.016)

20. Hassani MA, Durán P, Hacquard S. 2018 Microbial interactions within the plant holobiont. Microbiome 6, 58. (doi:10.1186/s40168-018-0445-0)

21. Liu M, Yu Z, Yu X, Xue Y, Huang B, Yang J. 2017 Invasion by Cordgrass Increases Microbial Diversity and Alters Community Composition in a Mangrove Nature Reserve. Frontiers in Microbiology 8. (doi:10.3389/fmicb.2017.02503)

22. Gu Y et al. 2016 Pathogen invasion indirectly changes the composition of soil microbiome via shifts in root exudation profile. Biology and Fertility of Soils 52, 997–1005. (doi:10.1007/s00374-016-1136-2)

23. Stringlis IA et al. 2018 MYB72-dependent coumarin exudation shapes root microbiome assembly to promote plant health. Proceedings of the National Academy of Sciences 115, E5213–E5222. (doi:10.1073/pnas.1722335115)

24. Vannier N, Agler M, Hacquard S. 2019 Microbiota-mediated disease resistance in plants. PLOS Pathogens 15, e1007740. (doi:10.1371/journal.ppat.1007740)

25. Rivett DW, Jones ML, Ramoneda J, Mombrikotb SB, Ransome E, Bell T. 2018 Elevated success of multispecies bacterial invasions impacts community composition during ecological succession. Ecology Letters 21, 516–524. (doi:10.1111/ele.12916)

26. Mallon CA, Le Roux X, van Doorn GS, Dini-Andreote F, Poly F, Salles JF. 2018 The impact of failure: unsuccessful bacterial invasions steer the soil microbial community away from the invader’s niche. The ISME Journal 12, 728–741. (doi:10.1038/s41396-017-0003-y)

27. Tilman D, Isbell F, Cowles JM. 2014 Biodiversity and Ecosystem Functioning. Annual Review of Ecology, Evolution, and Systematics 45, 471–493. (doi:10.1146/annurev-ecolsys-120213-091917)

28. Loper JE et al. 2012 Comparative Genomics of Plant-Associated Pseudomonas spp.: Insights into Diversity and Inheritance of Traits Involved in Multitrophic Interactions. PLoS Genetics 8, e1002784. (doi:10.1371/journal.pgen.1002784)

29. Manning P, van der Plas F, Soliveres S, Allan E, Maestre FT, Mace G, Whittingham MJ, Fischer M. 2018 Redefining ecosystem multifunctionality. Nature Ecology & Evolution 2, 427–436. (doi:10.1038/s41559-017-0461-7)

30. Hayward AC. 1991 Biology and Epidemiology of Bacterial Wilt Caused by Pseudomonas Solanacearum. Annual Review of Phytopathology 29, 65–87.

31. Jousset A, Schmid B, Scheu S, Eisenhauer N. 2011 Genotypic richness and dissimilarity opposingly affect ecosystem functioning. Ecology Letters 14, 537–545. (doi:10.1111/j.1461-0248.2011.01613.x)

32. Weidner S, Koller R, Latz E, Kowalchuk G, Bonkowski M, Scheu S, Jousset A. 2015 Bacterial diversity amplifies nutrient-based plant–soil feedbacks. Functional Ecology 29, 1341–1349. (doi:10.1111/1365-2435.12445)

33. Becker J, Eisenhauer N, Scheu S, Jousset A. 2012 Increasing antagonistic interactions cause bacterial communities to collapse at high diversity. Ecology Letters 15, 468–474. (doi:10.1111/j.1461-0248.2012.01759.x)

34. Jousset A, Schulz W, Scheu S, Eisenhauer N. 2011 Intraspecific genotypic richness and relatedness predict the invasibility of microbial communities. The ISME Journal 5, 1108–1114. (doi:10.1038/ismej.2011.9)

35. Soliveres S et al. 2016 Locally rare species influence grassland ecosystem multifunctionality. Philosophical Transactions of the Royal Society B: Biological Sciences 371, 20150269. (doi:10.1098/rstb.2015.0269)

36. Wei Z, Yang X, Yin S, Shen Q, Ran W, Xu Y. 2011 Efficacy of Bacillus-fortified organic fertiliser in controlling bacterial wilt of tomato in the field. Applied Soil Ecology 48, 152–159. (doi:doi.org/10.1016/j.apsoil.2011.03.013)

37. Cardenas E et al. 2010 Significant association between sulfate-reducing bacteria and uranium-reducing microbial communities as revealed by a combined massively parallel sequencing-indicator species approach. Applied and Environmental Microbiology 76, 6778–6786. (doi:10.1128/AEM.01097-10)

38. Caporaso JG et al. 2010 QIIME allows analysis of high-throughput community sequencing data. Nature methods 7, 335–336. (doi:10.1038/nmeth.f.303)

39. Edgar RC. 2013 UPARSE: highly accurate OTU sequences from microbial amplicon reads. Nature Methods 10, 996–998. (doi:10.1038/nmeth.2604)

40. Edgar RC, Haas BJ, Clemente JC, Quince C, Knight R. 2011 UCHIME improves sensitivity and speed of chimera detection. Bioinformatics 27, 2194–2200. (doi:10.1093/bioinformatics/btr381)

41. Margesin R, Płaza GA, Kasenbacher S. 2011 Characterization of bacterial communities at heavy-metal-contaminated sites. Chemosphere 82, 1583–1588. (doi:10.1016/j.chemosphere.2010.11.056)

42. Amato KR et al. 2013 Habitat degradation impacts black howler monkey (Alouatta pigra) gastrointestinal microbiomes. The ISME Journal 7, 1344–1353. (doi:10.1038/ismej.2013.16)

43. Cadotte MW, Davies TJ, Regetz J, Kembel SW, Cleland E, Oakley TH. 2010 Phylogenetic diversity metrics for ecological communities: integrating species richness, abundance and evolutionary history. Ecology Letters 13, 96–105. (doi:10.1111/j.1461-0248.2009.01405.x)

44. Doncaster CP. 2007 Structural Equation Modeling and Natural Systems. Fish and Fisheries 8, 368–369. (doi:10.1111/j.1467-2979.2007.00260.x)

45. Eisenhauer N, Bowker MA, Grace JB, Powell JR. 2015 From patterns to causal understanding: Structural equation modeling (SEM) in soil ecology. Pedobiologia 58, 65–72. (doi:10.1016/j.pedobi.2015.03.002)

46. Schmid B, Hector A, Saha P, Loreau M. 2008 Biodiversity effects and transgressive overyielding. Journal of Plant Ecology 1, 95–102. (doi:10.1093/jpe/rtn011)

47. Cleary DW, Bishop AH, Zhang L, Topp E, Wellington EMH, Gaze WH. 2016 Long-term antibiotic exposure in soil is associated with changes in microbial community structure and prevalence of class 1 integrons. FEMS Microbiology Ecology 92, fiw159. (doi:10.1093/femsec/fiw159)

48. Shade A, Klimowicz AK, Spear RN, Linske M, Donato JJ, Hogan CS, McManus PS, Handelsman J. 2013 Streptomycin Application Has No Detectable Effect on Bacterial Community Structure in Apple Orchard Soil. Applied and Environmental Microbiology 79, 6617–6625. (doi:10.1128/AEM.02017-13)

49. Freimoser FM, Pelludat C, Remus-Emsermann MNP. 2016 Tritagonist as a new term for uncharacterised microorganisms in environmental systems. The ISME Journal 10, 1–3. (doi:10.1038/ismej.2015.92)

50. Nelson WC, Stegen JC. 2015 The reduced genomes of Parcubacteria (OD1) contain signatures of a symbiotic lifestyle. Frontiers in Microbiology 6, 1–14. (doi:10.3389/fmicb.2015.00713)

51. Jousset A et al. 2017 Where less may be more: how the rare biosphere pulls ecosystems strings. The ISME Journal 11, 853–862. (doi:10.1038/ismej.2016.174)

52. Finkel OM et al. 2020 A single bacterial genus maintains root growth in a complex microbiome. Nature 587, 103–108. (doi:10.1038/s41586-020-2778-7)

53. Lee S-M, Kong HG, Song GC, Ryu C-M. 2021 Disruption of Firmicutes and Actinobacteria abundance in tomato rhizosphere causes the incidence of bacterial wilt disease. The ISME Journal, 330–347. (doi:10.1038/s41396-020-00785-x)

54. Li X, Jousset A, de Boer W, Carrión VJ, Zhang T, Wang X, Kuramae EE. 2019 Legacy of land use history determines reprogramming of plant physiology by soil microbiome. The ISME Journal 13, 738–751. (doi:10.1038/s41396-018-0300-0)

55. Wei Z, Gu Y, Friman V-P, Kowalchuk GA, Xu Y, Shen Q, Jousset A. 2019 Initial soil microbiome composition and functioning predetermine future plant health. Science Advances 5, eaaw0759. (doi:10.1126/sciadv.aaw0759)

