## Supplementary material for "Introduction of probiotic bacterial consortia promotes plant growth via impacts on the resident rhizosphere microbiome": Al supplementary Figures and Tables

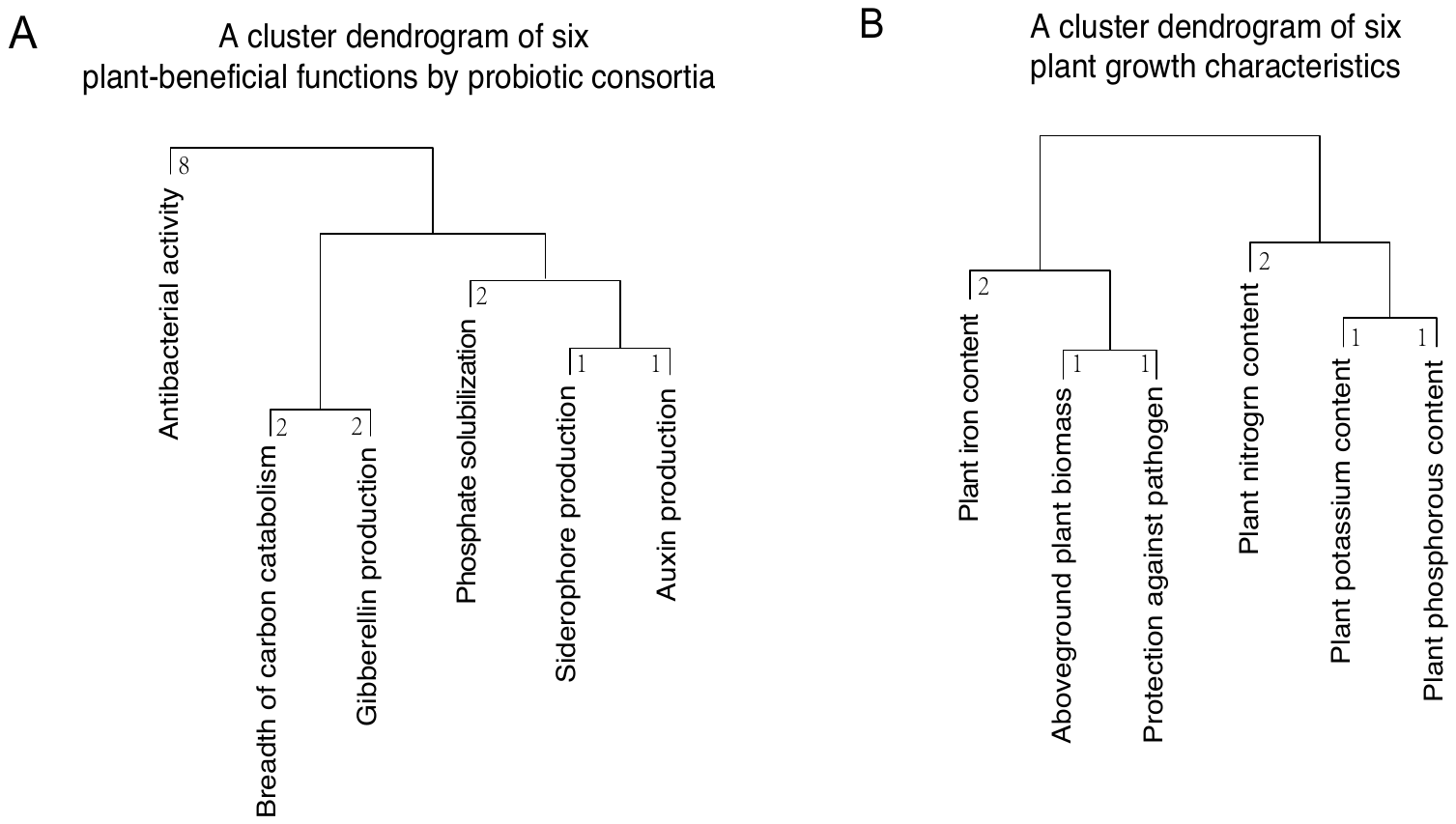

**Figure S1. Weighted cluster dendrograms of individual plant-beneficial functions by probiotic strains (A), and different plant growth characteristics (B) used for calculation of consortia multifunctionality and weighted average plant growth indexes.** A: A cluster dendrogram of six plant-beneficial functions used for calculation of consortia multifunctionality index. B: A cluster dendrogram of six plant growth characteristics used to calculate weighted average plant growth index. In both panels, the numbers at the tips of the dendrograms show the weighed values of each function or plant growth characteristic.

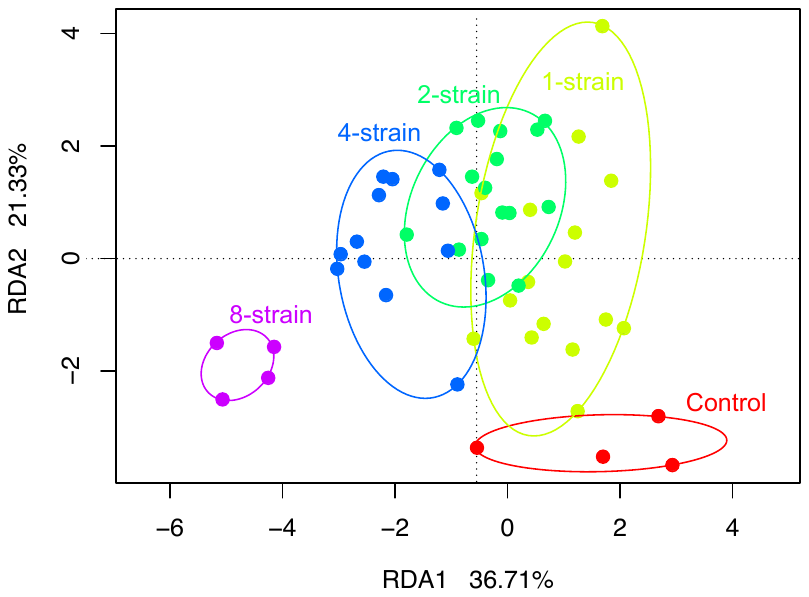

**Figure S2. Redundancy analysis (RDA) showing the impact of *Pseudomonas* consortium richness on the resident rhizosphere microbiome composition based on Amplicon Sequence Variant (ASV) table generated using DADA2 pipeline.** Ellipses enclosing points of each richness level show 95 % confidence intervals.

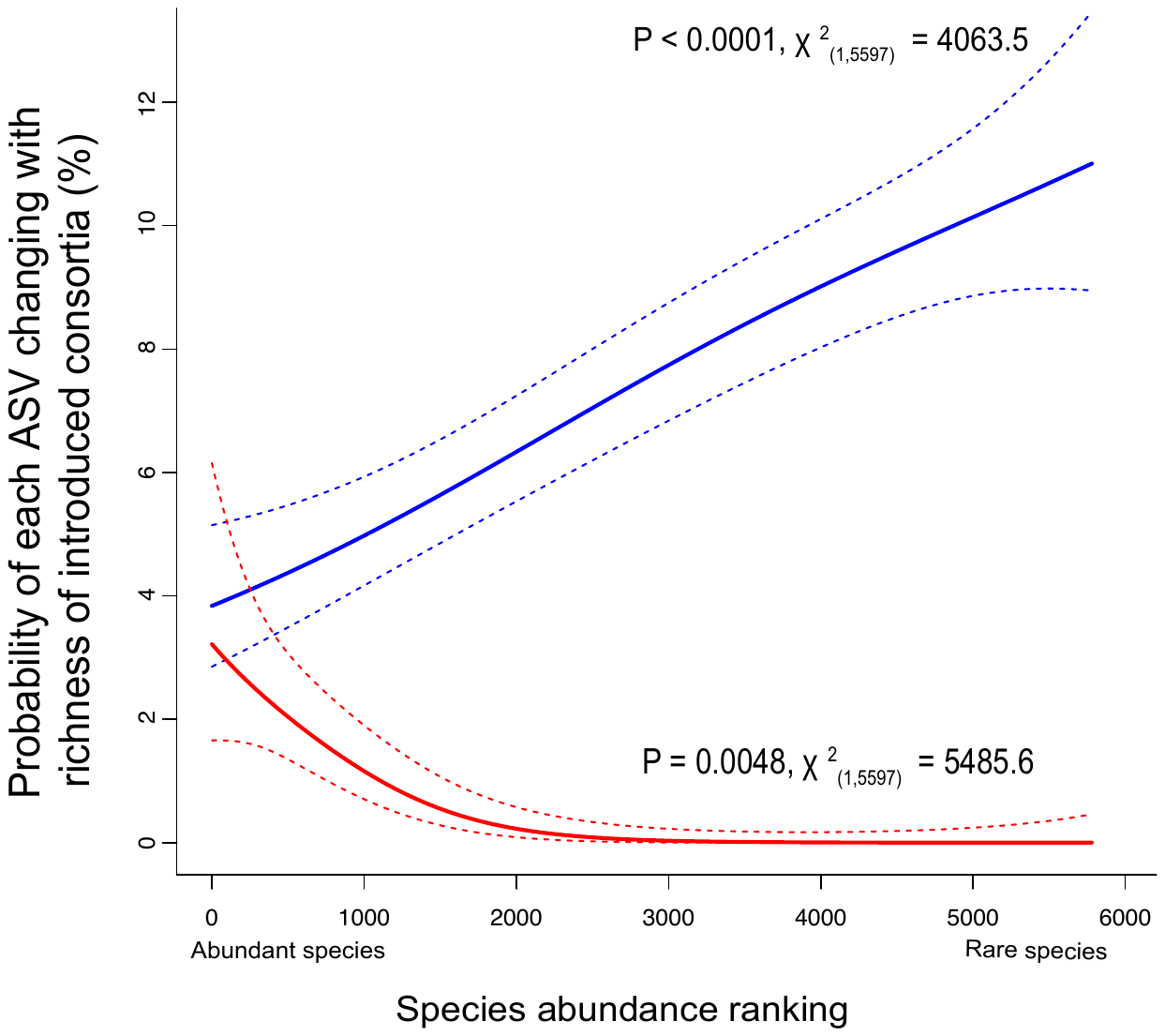

**Figure S3. Response of abundant and rare resident rhizosphere microbiome taxa to increasing richness of probiotic *Pseudomonas* consortia based on Amplicon Sequence Variant (ASV) table generated using DADA2 pipeline.** The curves show the relationship between ASV ranked abundance and the probability of the given ASV to significantly increase (blue line) or decrease (red line) along with *Pseudomonas* consortium richness. Dashed lines show 95% confidence intervals of binomial regressions.

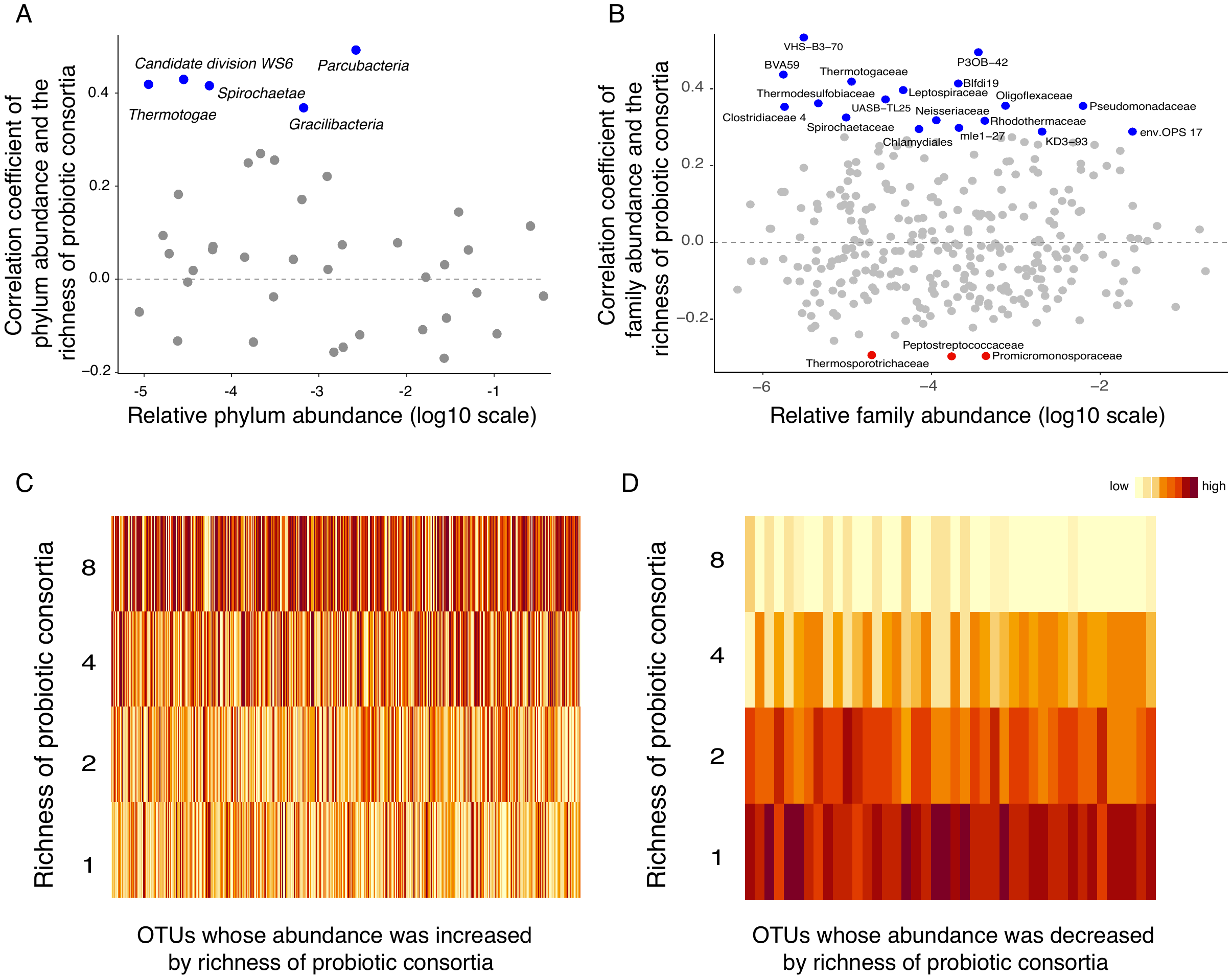

**Figure S4. Effect of probiotic *Pseudomonas* consortium richness on the abundance of different taxa present in the resident microbiome.** A-B: Relationship between correlation coefficients of taxa abundances along with consortium richness with relative taxa abundances at phylum (A) and family (B) levels. The blue and red dots indicate phyla and families whose relative abundances significantly increased or decreased, respectively, with the richness of probiotic consortia. Grey dots indicate phyla and families that showed non-significant changes. C: Heatmap of significantly increased OTU abundances along with the richness of probiotic consortia. D: Heatmap of significantly decreased OTU abundances along with the richness of probiotic consortia (in both panels, the yellow to red color gradient shows low to high relative OTU abundances in the rhizosphere microbiomes).

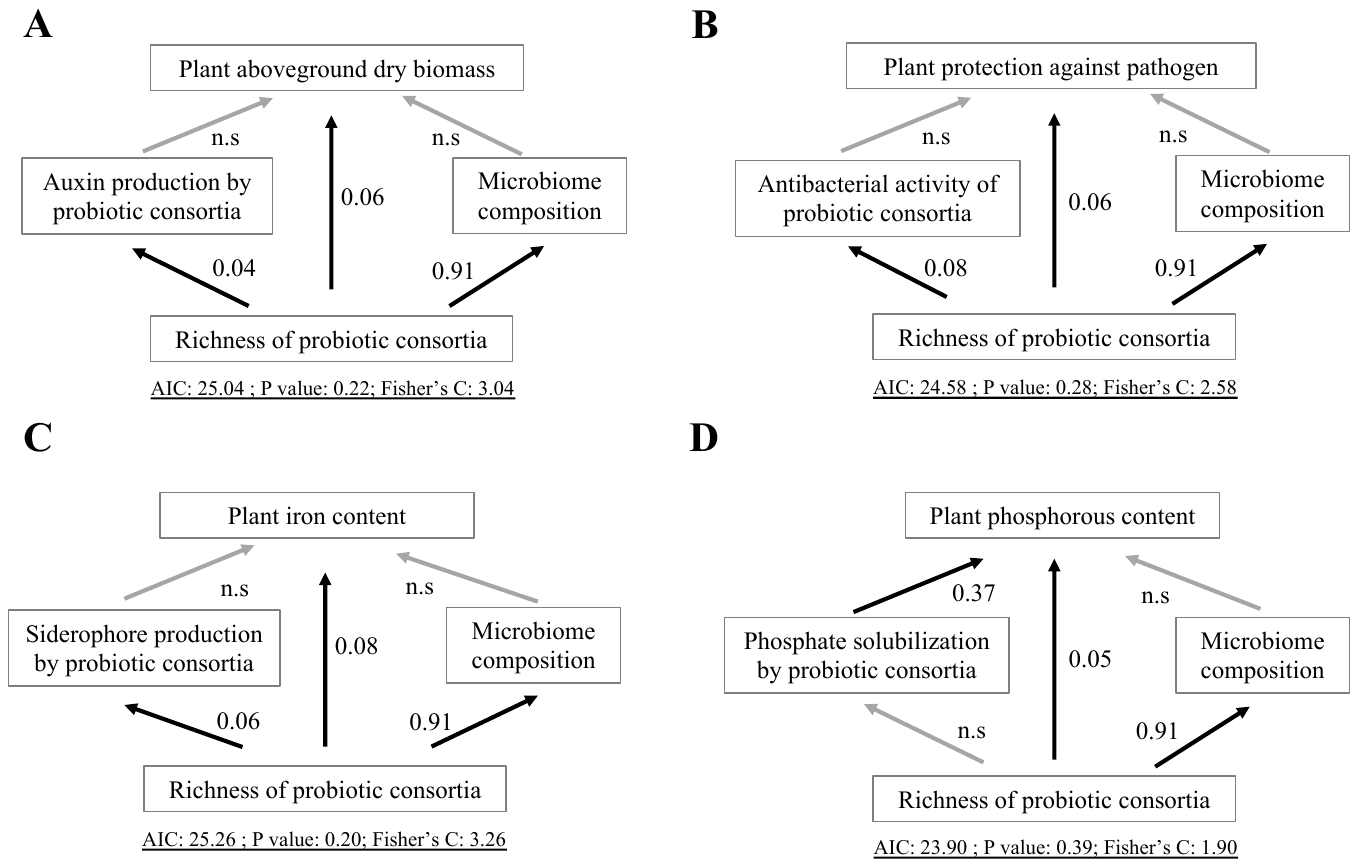

**Figure S5. Structural equation model (SEM) showing the direct and indirect relationships between the richness of probiotic *Pseudomonas* consortia, plant-beneficial functions of consortia measured *in vitro* and plant growth characteristics.** For each trait of interest (Panels A-D), the impact of consortium richness was evaluated as its direct effect (expression of the underlying traits *in vitro*; left middle boxes) and indirect effect (mediated by changes in the resident microbiome composition, RDA1; right middle boxes). Panels A to D show the effect of probiotic consortium richness on aboveground dry plant biomass, plant protection against pathogen infection, plant iron content, and plant phosphorus content, respectively. In each panel, black and grey arrows indicate significantly positive or non-significant relationships between different variables, respectively. The numbers beside arrows show standardized correlation coefficients (relative effect sizes of non-significant correlations are not shown) and the parameters below each panel show the robustness of the SEM model. SEM explaining plant potassium and nitrogen contents were not included because of all relationships were non-significant.

**Table S1.** **List of *Pseudomonas* strains used for assembling introduced bacterial consortia in this study.**

| *Pseudomonas* ID | *Pseudomonas* strain | Origin | Reference |
| --- | --- | --- | --- |
| CHA0 | *Pseudomonas protegens* | Tobacco, Switzerland | Natsch A. et al. ,Appl Environ Microbiol 60:2553-2560, 1994 |
| F113 | *Pseudomonas kilonensis* | Sugar beet, Ireland | Shanahan P. et al. ,Appl Environ Microbiol 58:353-358, 1992 |
| Phl1c2 | *Pseudomonas fluorescens* | Tomato, France | De La Fuente L. et al. , FEMS Microbiol Ecol 56:64-78, 2006 |
| Pf-5 | *Pseudomonas protegens* | Cotton, USA | Howell C.R. et al. , Phytopathology 69:480-482,1979 |
| Q2-87 | *Pseudomonas fluorescens* | Wheat, USA | Bangera M.G. et al. , J Bacteriol 181:3155-3163,1999 |
| Q8r1-96 | *Pseudomonas brassicacearum* | Wheat, USA | Raaijmakers J.M. et al. , Mol Plant Microbe Interact 11:144-152,1998 |
| 1m1-96 | *Pseudomonas fluorescens* | Wheat, USA | Raaijmakers J.M. et al. , Appl Environ Microbiol 67:2545-2554,2001 |
| mvp1-4 | *Pseudomonas fluorescens* | Pea, USA | Landa BB et al. , Appl Environ Microbiol 68:3226-3237,2002 |

Note: *Pseudomonas spp.* strains contained a marker gene *PhlD*, which was otherwise found at low abundances in the non-sterile agricultural field soil used for our experiments, allowing the quantification of total inoculant bacterial densities during the experiment. See also (Loper JE, Hassan KA, Mavrodi DV, Davis EW, 2nd, Lim CK, Shaffer BT, Elbourne LD, Stockwell VO, Hartney SL, Breakwell K, Henkels MD, Tetu SG, Rangel LI, Kidarsa TA, Wilson NL, van de Mortel JE, Song C, Blumhagen R, Radune D, Hostetler JB, Brinkac LM, Durkin AS, Kluepfel DA, Wechter WP, Anderson AJ, Kim YC, Pierson LS, 3rd, Pierson EA, Lindow SE, Kobayashi DY, Raaijmakers JM, Weller DM, Thomashow LS, Allen AE, Paulsen IT., vol 8, p e1002784,2012 )for an updated classiﬁcation of the Pseudomonas strains; + toxin production, DAPG 2, 4-diacetylphloroglucinol, PLT pyoluteorin, PRN pyrrolnitrin, HCN hydrogen cyanide, AprA extracellular protease.

**Table S2.** **Composition of the probiotic *Pseudomonas* consortia used in this study.**

| *Pseudomonas* consortium ID | *Pseudomonas* strain | | | | | | | | Consortium richness |
| --- | --- | --- | --- | --- | --- | --- | --- | --- | --- |
|  | mvp1-4 | Q2-87 | CHA0 | F113 | Phl1c2 | Pf-5 | 1m1-96 | Q8r1-96 |  |
| 1 | 1 | 0 | 0 | 0 | 0 | 0 | 0 | 0 | 1 |
| 2 | 0 | 1 | 0 | 0 | 0 | 0 | 0 | 0 | 1 |
| 3 | 0 | 0 | 1 | 0 | 0 | 0 | 0 | 0 | 1 |
| 4 | 0 | 0 | 0 | 1 | 0 | 0 | 0 | 0 | 1 |
| 5 | 0 | 0 | 0 | 0 | 1 | 0 | 0 | 0 | 1 |
| 6 | 0 | 0 | 0 | 0 | 0 | 1 | 0 | 0 | 1 |
| 7 | 0 | 0 | 0 | 0 | 0 | 0 | 1 | 0 | 1 |
| 8 | 0 | 0 | 0 | 0 | 0 | 0 | 0 | 1 | 1 |
| 9 | 1 | 0 | 0 | 0 | 0 | 0 | 0 | 0 | 1 |
| 10 | 0 | 1 | 0 | 0 | 0 | 0 | 0 | 0 | 1 |
| 11 | 0 | 0 | 1 | 0 | 0 | 0 | 0 | 0 | 1 |
| 12 | 0 | 0 | 0 | 1 | 0 | 0 | 0 | 0 | 1 |
| 13 | 0 | 0 | 0 | 0 | 1 | 0 | 0 | 0 | 1 |
| 14 | 0 | 0 | 0 | 0 | 0 | 1 | 0 | 0 | 1 |
| 15 | 0 | 0 | 0 | 0 | 0 | 0 | 1 | 0 | 1 |
| 16 | 0 | 0 | 0 | 0 | 0 | 0 | 0 | 1 | 1 |
| 17 | 1 | 1 | 0 | 0 | 0 | 0 | 0 | 0 | 2 |
| 18 | 0 | 1 | 1 | 0 | 0 | 0 | 0 | 0 | 2 |
| 19 | 1 | 0 | 0 | 1 | 0 | 0 | 0 | 0 | 2 |
| 20 | 0 | 1 | 0 | 0 | 1 | 0 | 0 | 0 | 2 |
| 21 | 1 | 0 | 0 | 0 | 0 | 0 | 1 | 0 | 2 |
| 22 | 0 | 0 | 1 | 1 | 0 | 0 | 0 | 0 | 2 |
| 23 | 1 | 0 | 0 | 0 | 0 | 0 | 0 | 1 | 2 |
| 24 | 0 | 0 | 1 | 0 | 0 | 1 | 0 | 0 | 2 |
| 25 | 0 | 0 | 0 | 1 | 1 | 0 | 0 | 0 | 2 |
| 26 | 0 | 0 | 1 | 0 | 0 | 0 | 1 | 0 | 2 |
| 27 | 0 | 1 | 0 | 0 | 0 | 1 | 0 | 0 | 2 |
| 28 | 0 | 0 | 0 | 1 | 0 | 0 | 0 | 1 | 2 |
| 29 | 0 | 0 | 0 | 0 | 1 | 0 | 0 | 1 | 2 |
| 30 | 0 | 0 | 0 | 0 | 1 | 1 | 0 | 0 | 2 |
| 31 | 0 | 0 | 0 | 0 | 0 | 1 | 1 | 0 | 2 |
| 32 | 0 | 0 | 0 | 0 | 0 | 0 | 1 | 1 | 2 |
| 33 | 1 | 1 | 0 | 1 | 1 | 0 | 0 | 0 | 4 |
| 34 | 0 | 1 | 0 | 1 | 0 | 1 | 0 | 1 | 4 |
| 35 | 1 | 0 | 0 | 1 | 1 | 1 | 0 | 0 | 4 |
| 36 | 0 | 1 | 0 | 0 | 1 | 0 | 1 | 1 | 4 |
| 37 | 1 | 1 | 1 | 1 | 0 | 0 | 0 | 0 | 4 |
| 38 | 0 | 1 | 0 | 0 | 0 | 1 | 1 | 1 | 4 |
| 39 | 1 | 0 | 0 | 0 | 1 | 1 | 1 | 0 | 4 |
| 40 | 0 | 1 | 1 | 0 | 1 | 0 | 0 | 1 | 4 |
| 41 | 1 | 0 | 1 | 0 | 0 | 1 | 1 | 0 | 4 |
| 42 | 0 | 0 | 1 | 1 | 0 | 1 | 0 | 1 | 4 |
| 43 | 1 | 0 | 1 | 1 | 0 | 0 | 1 | 0 | 4 |
| 44 | 0 | 0 | 1 | 0 | 1 | 0 | 1 | 1 | 4 |
| 45 | 1 | 1 | 1 | 1 | 1 | 1 | 1 | 1 | 8 |
| 46 | 1 | 1 | 1 | 1 | 1 | 1 | 1 | 1 | 8 |
| 47 | 1 | 1 | 1 | 1 | 1 | 1 | 1 | 1 | 8 |
| 48 | 1 | 1 | 1 | 1 | 1 | 1 | 1 | 1 | 8 |

All *Pseudomonas* monocultures were replicated twice except for 8-species communities that were replicated four times. In other diversity levels, each *Pseudomonas* strain was included equally often into assembled communities. In the table, 1 denotes for the presence and 0 for the absence of the strain in the given community. This experimental design is not fully factorial. Instead, it aims to keep the total abundance of bacterial strains the same in every consortium and ensuring that each strain is present equally often at each richness level, thereby allowing separation of richness and strain identity effects from each other.

**Table S3.** **List of plant-beneficial bacterial functions and their effects on the plant**

| **Plant beneficial function** | **Effect on the plant** | **References** |
| --- | --- | --- |
| Breadth of carbon catabolism | Establishment in the rhizosphere; pathogen inhibition via resource competition | (Hu et al., 2016) |
| Antibiotic production | Pathogen direct inhibition | (Jousset et al., 2011) |
| Gibberellin production | Improvement of plant growth | (Bottini et al., 2004) |
| Auxin production | Improvement of plant growth | (Zhao, 2010) |
| Phosphate solubilization | Improvement of plant phosphorus uptake | (Shen et al., 2011) |
| Siderophore production | Facilitation of plant iron uptake | (Ahmed and Holmström, 2014) |

Ahmed E, Holmström SJM. 2014. Siderophores in environmental research: roles and applications: Siderophores in environmental research. *Microbial Biotechnology* **7**:196–208. doi:10.1111/1751-7915.12117

Bottini R, Cassán F, Piccoli P. 2004. Gibberellin production by bacteria and its involvement in plant growth promotion and yield increase. *Applied Microbiology and Biotechnology* **65**. doi:10.1007/s00253-004-1696-1

Hu J, Wei Z, Friman V-P, Gu S, Wang X, Eisenhauer N, Yang T, Ma J, Shen Q, Xu Y, Jousset A. 2016. Probiotic diversity enhances rhizosphere microbiome function and plant disease suppression. *mBio* **7**:e01790-16. doi:10.1128/mBio.01790-16

Jousset A, Rochat L, Lanoue A, Bonkowski M, Keel C, Scheu S. 2011. Plants Respond to Pathogen Infection by Enhancing the Antifungal Gene Expression of Root-Associated Bacteria. *Molecular Plant-Microbe Interactions* **24**:352–358. doi:10.1094/MPMI-09-10-0208

Shen J, Yuan L, Zhang J, Li H, Bai Z, Chen X, Zhang W, Zhang F. 2011. Phosphorus Dynamics: From Soil to Plant. *Plant Physiology* **156**:997–1005. doi:10.1104/pp.111.175232

Zhao Y. 2010. Auxin Biosynthesis and Its Role in Plant Development. *Annual Review of Plant Biology* **61**:49–64. doi:10.1146/annurev-arplant-042809-112308

**Table S4. Origin of the data used in this paper**

| Traits | Data | Origin |
| --- | --- | --- |
| *Pseudomonas* *in*  *vitro* traits | Carbon utilization | Hu et al, mBio, 2016 |
|  | Antibacterial activity |  |
| Plant traits | Plant disease incidence |  |
| *Pseudomonas* *in*  *vitro* traits | Gibberellin production | Hu et al, SBB, 2017 |
|  | Auxin production |  |
|  | Phosphate solubilization |  |
|  | Siderophore production |  |
| Plant traits | Aboveground dry plant biomass |  |
|  | Plant potassium content |  |
|  | Plant phosphorous content |  |
|  | Plant iron content |  |
|  | Plant nitrogen content |  |
| Rhizosphere bacterial data | *PhlD* gene quantification |  |
|  | Bacterial 16S rRNA gene quantification |  |
| Rhizosphere bacterial data | 16S rRNA amplicon sequence-based microbiome analysis | This study; rhizosphere samples originated from the greenhouse experiment described in Hu et al, SBB, 2017 |

**Table S5.** ***Pseudomonas* strain identity and consortium richness effects on the plant-beneficial functions (measured *in vitro*) and plant growth characteristics (measured *in vivo*).**

|  | Introduced *Pseudomonas* strain identity | | | | | | | | | |
| --- | --- | --- | --- | --- | --- | --- | --- | --- | --- | --- |
|  |  | mvp1-4 | Q2-87 | CHA0 | F113 | Phl1c2 | Pf-5 | 1m1-96 | Q8r1-96 | Consortium  richness |
| Consortium multifunctionality index | F | 1.76 | 2.06 | 3.54 | 1.57 | 1.84 | 8.29 | 1.26 | 1.01 | 1263.01 |
|  | P | 0.19 | 0.16 | 0.07 | 0.22 | 0.18 | **0.01** | 0.27 | 0.32 | **<0.0001** |
| Breadth of carbon catabolism | F | 1.79 | 1.22 | 1.49 | 1.43 | 2.52 | 2.83 | 2.22 | 1.38 | 4737.87 |
|  | P | 0.19 | 0.28 | 0.23 | 0.24 | 0.12 | 0.10 | 0.14 | 0.25 | **<0.0001** |
| Siderophore production | F | 1.64 | 2.56 | 2.17 | 0.69 | 1.32 | 6.81 | 3.30 | 1.59 | 299.32 |
|  | P | 0.21 | 0.12 | 0.15 | 0.41 | 0.26 | **0.01** | 0.08 | 0.21 | **<0.0001** |
| Auxin production | F | 0.97 | 0.85 | 3.46 | 1.90 | 3.89 | 1.66 | 2.19 | 2.07 | 417.09 |
|  | P | 0.33 | 0.36 | 0.07 | 0.18 | 0.06 | 0.20 | 0.15 | 0.16 | **<0.0001** |
| Phosphate solubilization | F | 0.09 | 1.05 | 4.02 | 2.42 | 4.52 | 2.79 | **0.03** | 0.42 | 417.09 |
|  | P | 0.77 | 0.31 | **0.05** | 0.13 | **0.04** | 0.10 | 0.86 | 0.52 | **<0.0001** |
| Gibberellin production | F | 1.23 | 1.64 | 1.48 | 1.13 | 1.40 | 1.36 | 1.60 | 1.47 | 9299.27 |
|  | P | 0.27 | 0.21 | 0.23 | 0.29 | 0.24 | 0.25 | 0.21 | 0.23 | **<0.0001** |
| Antibacterial activity | F | 3.36 | 3.18 | 5.59 | 1.48 | 0.79 | 23.04 | 1.12 | 0.65 | 241.61 |
|  | P | 0.07 | 0.08 | **0.02** | 0.23 | 0.38 | **<0.0001** | 0.30 | 0.42 | **<0.0001** |
| Weighted average of plant growth index | F | 1.75 | 2.24 | 1.64 | 2.24 | 1.34 | 1.68 | 1.52 | 2.05 | 314.64 |
|  | P | 0.19 | 0.14 | 0.21 | 0.14 | 0.25 | 0.20 | 0.22 | 0.16 | **<0.0001** |
| Aboveground plant dry biomass | F | 3.57 | 0.87 | 3.54 | 2.53 | 3.21 | 1.37 | 6.41 | 1.47 | 339.14 |
|  | P | 0.07 | 0.36 | 0.07 | 0.12 | 0.08 | 0.25 | **0.02** | 0.23 | **<0.0001** |
| Plant potassium content | F | 1.96 | 3.23 | 1.15 | 1.47 | 0.51 | 3.61 | 0.82 | 1.53 | 56.62 |
|  | P | 0.17 | 0.08 | 0.29 | 0.23 | 0.48 | 0.06 | 0.37 | 0.22 | **<0.0001** |
| Plant phosphorus content | F | 0.42 | 1.83 | 7.37 | 1.53 | 2.05 | 3.32 | 2.24 | 1.06 | 71.60 |
|  | P | 0.52 | 0.18 | **0.01** | 0.22 | 0.16 | 0.08 | 0.14 | 0.31 | **<0.0001** |
| Plant iron content | F | 5.71 | 1.00 | 2.47 | 6.37 | 1.29 | 6.21 | 6.11 | 3.56 | 24.95 |
|  | P | **0.02** | 0.32 | 0.12 | **0.02** | 0.26 | 0.02 | **0.02** | 0.07 | **<0.0001** |
| Plant nitrogen content | F | 1.69 | 1.21 | 0.10 | 1.54 | 0.86 | 0.23 | 0.13 | 1.48 | 69.00 |
|  | P | 0.20 | 0.28 | 0.75 | 0.22 | 0.36 | 0.63 | 0.72 | 0.23 | **<0.0001** |
| Protection against pathogen infection | F | 0.62 | 3.20 | 1.40 | 2.85 | 0.62 | 1.87 | 2.39 | 3.33 | 336.15 |
|  | P | 0.44 | 0.08 | 0.24 | 0.10 | 0.44 | 0.18 | 0.13 | 0.08 | **<0.0001** |
|  | df | 1 | 1 | 1 | 1 | 1 | 1 | 1 | 1 | 1 |
|  | No. of residuals: 40 | | |  |  |  |  |  |  |  |

F denotes for variance ratio, P for error probability, and significant results (p<0.05) are highlighted in bold.

**Table S6. *Pseudomonas* strain identity effects on the ratio of *phlD*/16S rRNA gene copy numbers and the total bacterial abundances in the rhizosphere.**

|  |  | *Pseudomonas* strain identity | | | | | | | |  |
| --- | --- | --- | --- | --- | --- | --- | --- | --- | --- | --- |
|  |  | mvp1-4 | Q2-87 | CHA0 | F113 | Phl1c2 | Pf-5 | 1m1-96 | Q8r1-96 | Consortium richness |
| The ratio of *phlD*/16S rRNA gene copy numbers in the rhizosphere | F | 4.30 | 3.36 | 3.73 | 1.02 | 1.36 | 2.60 | 0.90 | 1.86 | 4301.8 |
|  | P | **0.04** | 0.07 | 0.06 | 0.32 | 0.25 | 0.11 | 0.35 | 0.18 | **<0.0001** |
| Total bacterial abundances in the rhizosphere | F | 1.18 | 1.34 | 1.29 | 1.20 | 1.29 | 1.26 | 1.21 | 1.21 | 2110.4 |
|  | P | 0.28 | 0.25 | 0.26 | 0.28 | 0.26 | 0.27 | 0.28 | 0.28 | **<0.0001** |
|  | df | 1 | 1 | 1 | 1 | 1 | 1 | 1 | 1 |  |
|  | No. of residuals: 40 | | | | | | | | |  |

F denotes for variance ratio, P for error probability, and significant results (p<0.05) are highlighted in bold.

**Table S7.** ***Pseudomonas* strain identity effect on resident rhizosphere microbiome diversity.**

| Resident rhizosphere microbiome diversity index |  | *Pseudomonas* strain identity | | | | | | | |  |
| --- | --- | --- | --- | --- | --- | --- | --- | --- | --- | --- |
|  |  | mvp1-4 | Q2-87 | CHA0 | F113 | Phl1c2 | Pf-5 | 1m1-96 | Q8r1-96 | Consortium richness |
| Community phylogenetic  abundance evenness | F | 1.63 | 0.98 | 1.73 | 0.91 | 1.28 | 1.26 | 1.52 | 1.57 | 4301.8 |
|  | P | 0.21 | 0.33 | 0.20 | 0.34 | 0.26 | 0.27 | 0.23 | 0.22 | **<0.0001** |
| Community phylogenetic diversity | F | 1.70 | 1.30 | 1.49 | 0.98 | 1.32 | 1.19 | 1.44 | 1.37 | 2110.4 |
|  | P | 0.20 | 0.26 | 0.23 | 0.33 | 0.26 | 0.28 | 0.24 | 0.25 | **<0.0001** |
| Community OTU richness | F | 1.75 | 1.25 | 1.31 | 0.95 | 1.34 | 1.16 | 1.39 | 1.43 | 1191.1 |
|  | P | 0.19 | 0.27 | 0.26 | 0.34 | 0.25 | 0.29 | 0.24 | 0.24 | **<0.0001** |
| Community Shannon diversity | F | 1.42 | 1.15 | 1.52 | 1.09 | 1.21 | 1.22 | 1.33 | 1.29 | 3631.1 |
|  | P | 0.24 | 0.29 | 0.23 | 0.30 | 0.28 | 0.28 | 0.26 | 0.26 | **<0.0001** |
| Community Pielou evenness | F | 1.34 | 1.16 | 1.52 | 1.11 | 1.20 | 1.24 | 1.31 | 1.26 | 4986.44 |
|  | P | 0.25 | 0.29 | 0.23 | 0.30 | 0.28 | 0.27 | 0.26 | 0.27 | **<0.0001** |
|  | df | 1 | 1 | 1 | 1 | 1 | 1 | 1 | 1 |  |
|  | No. of residuals: 40 | | | | | | | | |  |

F denotes for variance ratio, P for error probability, and significant results (p<0.05) are highlighted in bold.

**Table S8.** **List of bacterial phyla that were statistically significantly correlated with probiotic consortium richness.**

| Phylum name | P value | Correlation coefficient |
| --- | --- | --- |
| Parcubacteria | 0.0004 | 0.49 |
| Candidate.division.WS6 | 0.0024 | 0.43 |
| Thermotogae | 0.0031 | 0.42 |
| Spirochaetae | 0.0033 | 0.42 |
| Gracilibacteria | 0.0101 | 0.37 |

**Table S9.** **Information of resident microbiome OTUs that significantly correlated with probiotic consortium OTUs within modules 1 and 2 of the co-occurrence network.** Blue colors indicate those OTUs that were positively correlated with the richness of the probiotic consortia, green colors indicate those OTUs belonging to the genus *Pseudomonas* that increased with the richness of the probiotic consortia and grey colors indicate those OTUs that had non-significant correlation with the richness of introduced consortia.

|  | **OTU name** | **Color in the network** | **Phylum name** | **Class name** | **Order name** | **Family name** | **Genus name** |
| --- | --- | --- | --- | --- | --- | --- | --- |
| Module 1 | OTU8735 | grey | Actinobacteria | Actinobacteria | Acidimicrobiales | uncultured | uncultured |
|  | OTU3936 | blue | Proteobacteria | Alphaproteobacteria | Caulobacterales | Caulobacteraceae | uncultured |
|  | OTU5145 | blue | Saccharibacteria | norank | norank | norank | norank |
|  | OTU7663 | blue | Actinobacteria | Actinobacteria | Acidimicrobiales | Acidimicrobiaceae | Illumatobacter |
|  | OTU1926 | blue | Bacteroidetes | Sphingobacteriia | Sphingobacteriales | NS11-12 marine group | norank |
|  | OTU4429 | blue | Parcubacteria | norank | norank | norank | norank |
|  | OTU663 | blue | Actinobacteria | Actinobacteria | Acidimicrobiales | Acidimicrobiales Incertae Sedis | Candidatus Microthrix |
|  | OTU6750 | blue | Bacteroidetes | Sphingobacteriia | Sphingobacteriales | Sphingobacteriaceae | Mucilaginibacter |
|  | OTU5266 | blue | Proteobacteria | Gammaproteobacteria | 34P16 | norank | norank |
|  | OTU2183 | blue | Parcubacteria | norank | norank | norank | norank |
|  | OTU1369 | blue | Proteobacteria | Gammaproteobacteria | Legionellales | Coxiellaceae | Aquicella |
|  | OTU8653 | blue | Proteobacteria | Alphaproteobacteria | Rhodospirillales | Rhodospirillaceae | Dongia |
|  | OTU8407 | blue | Actinobacteria | Actinobacteria | Streptosporangiales | Thermomonosporaceae | Thermomonospora |
|  | OTU498 | blue | Actinobacteria | Actinobacteria | Micrococcales | Demequinaceae | Demequina |
|  | OTU8478 | green | Proteobacteria | Gammaproteobacteria | Pseudomonadales | Pseudomonadaceae | Pseudomonas |
|  | OTU447 | blue | Proteobacteria | Gammaproteobacteria | Xanthomonadales | Xanthomonadaceae | uncultured |
|  | OTU398 | blue | Proteobacteria | Gammaproteobacteria | Xanthomonadales | uncultured | uncultured |
|  | OTU5111 | blue | Bacteroidetes | Cytophagia | Order II | Rhodothermaceae | uncultured |
| Module 2 | OTU2720 | blue | Proteobacteria | Deltaproteobacteria | Myxococcales | Blfdi19 | norank |
|  | OTU6162 | blue | Proteobacteria | Gammaproteobacteria | Xanthomonadales | Xanthomonadaceae | Luteimonas |
|  | OTU4923 | green | Proteobacteria | Gammaproteobacteria | Pseudomonadales | Pseudomonadaceae | Pseudomonas |
|  | OTU2574 | blue | Proteobacteria | Betaproteobacteria | Burkholderiales | Oxalobacteraceae | Paucimonas |

**Table S10. The effect of in vitro plant-beneficial functions measured at the consortium level in explaining the variation in the resident rhizosphere microbiome composition.**

| Plant-beneficial function measured at the consortium level | | Resident microbiome composition | | | | | |
| --- | --- | --- | --- | --- | --- | --- | --- |
|  |  | Principal component 1 | | | Principal component 2 | | |
|  |  | df | F | P | df | F | P |
| Competition | Breadth of carbon catabolism |  | n.r | n.r | 1 | 31.62 | **<0.0001** |
|  | Antibacterial activity | 1 | 2.33 | 0.1342 |  | n.r | n.r |
| Phytohormone production | Auxin  production | 1 | 14.97 | **0.0003** | 1 | 4.81 | **0.0335** |
|  | Gibberellin production | 1 | 21.61 | **<0.0001** |  | n.r | n.r |
| Nutrient availability | Phosphate solubilization |  | n.r | n.r |  | n.r | n.r |
|  | Siderophore production |  | n.r | n.r |  | n.r | n.r |
| Model summary | | 44 | R^2^ = 0.47  P < 0.0001 | | 45 | R^2^ = 0.42  P < 0.0001 | |

Stepwise general linear model was used for explaining the variation of resident rhizosphere microbiome composition based on *in vitro* plant-beneficial functions measured at the consortium level. Significant results (p < 0.05) are highlighted in bold and “n.r” denotes for a variable that was not retained in the final model.
