## Supplementary materials and methods for "Introduction of probiotic bacterial consortia promotes plant growth via impacts on the resident rhizosphere microbiome"

**Supplementary material and methods**

***Niche breadth.*** We assessed the resource use of the eight *Pseudomonas spp.* strains on 48 different single carbon resources representative of tomato root exudates following an established procedures [1]*.* Briefly, all bacteria were grown for 48 h (initial OD_600_ = 0.05) at 28 °C in Ornston and Stanier (OS) [2]) minimal medium supplemented with 10 mM of the target resource on 96-well microplates [1]*.* Wells with an OD_600_ > 0.5 (Spectra Max M5 Plate reader, Molecular Devices, Sunnyvale, CA, USA) were scored as positive for bacterial growth. Measurements were replicated three times for each strain for all carbon resources. The niche breadth estimate of all consortia was calculated as the number of resources consumed by at least one of the community members based on monoculture measurements [3].

***Siderophore production.*** We grew all *Pseudomonas* monocultures and consortia in MKB medium [4] for 48 h at 30 °C with shaking (170 rpm). After centrifugation (10’000 g, 5 min), siderophore concentration in the supernatant was measured with a CAS-shuttle assay [5]. Siderophore concentration of the supernatant was calculated using the formula developed by Swapan et al [5] and all measurements were replicated for three times for each monoculture and consortium.

***Auxin production.*** We grew all *Pseudomonas* monocultures and consortia in Landy’s medium [6] for 72 h at 22 °C in the dark with shaking (90 rpm). Bacterial cultures were then centrifuged (10’000 g, 5 min) and auxin concentration in the supernatants (ng/mL) determined with the IAA|Auxin ELISA Quantitation Kit (R&D, Shanghai, China) following the manufacturer’s protocol. Measurements were replicated for three times for each monoculture and consortium.

***Gibberellin production.*** We grew all *Pseudomonas* monocultures and consortia in nutrient broth for 72 h at 30 °C in the dark with shaking (200 rpm). After centrifugation (10’000 g for 5 min), gibberellin concentration was measured from the bacterial supernatant (pmol/L) using the Gibberellic Acid ELISA Quantitation Kit (R&D, Shanghai, China) according to the manufacturer's protocol. Measurements were replicated for three times for each monoculture and consortium.

***Phosphate solubilization.*** We grew all *Pseudomonas* monocultures and consortia in NBRIP medium [7] for 7 days at 30 °C with shaking (170 rpm). After centrifugation (10’000 g, 10 min) the soluble phosphate concentration (μg/mL) in the supernatant was measured using the molybdenum antimony colorimetric method [8]. Measurements were replicated for three times for each monoculture and consortium.

***Antibacterial activity against plant pathogenic R. solanacearum bacterium.*** We grew all *Pseudomonas* monocultures and consortia in nutrient broth for 30 h (30 °C, 170 rpm), after cells were pelleted by centrifugation (4000 g, 3 min), and inhibition experiments started immediately. Briefly, 20 μL of cell-free supernatant was added to a fresh *R. solanacearum* QL-Rs1115 pathogen culture (180 µL, OD_600_ = 0.05) in M-SMSA medium [9]. Control treatments received 20 µL nutrient broth medium supernatant free of *Pseudomonas* secondary metabolites. Bacteria were grown for 24 h (30 °C, 170 rpm) before measuring densities as optical density at 600 nm using a spectrophotometer (Spectra Max M5 Plate reader, Molecular Devices, Sunnyvale, CA, USA). Antibacterial activity was defined as the percentage pathogen growth reduction by supernatant compared to pathogen growth in the control treatment [3]. Measurements were replicated for three times for each monoculture and consortium.

***Plant growth-promotion experiment.*** The soil used in both greenhouse experiment was a yellow-brown earth (Udic Argosol), sieved at 5 mm, homogenized throughout and quantified to have following chemical characteristics: pH of 5.8, NH_4_^+^ of 2.01 mg/kg, NO_3_^-^ of 22.98 mg/kg, organic matter content of 24.00 g/kg, water soluble organic carbon of 101.92 mg/kg, water soluble organic nitrogen of 66.55 mg/kg, available phosphorus of 173.1 mg/kg, available potassium of 178 mg/kg, total N of 0.23% and total C of 2.1%). No additional fertilizer was applied in these two greenhouse experiments as pilot experiments revealed that the soil already contained enough nutrients to support plant growth.

After 10 days of growth in seedling trays, 48 trays were inoculated with each *Pseudomonas spp.* consortium following the substitutive design described earlier (Table S2) using the root drenching method with a final concentration of 5.0 × 10^7^ CFU of bacteria g^-1^ soil [10]. The remaining 4 trays were used for *Pseudomonas-*free control treatment. Tomato plants were grown for 60 days in a greenhouse (natural temperature variation ranging from 25 °C to 35 °C), watered regularly with sterile water and trays rearranged randomly every two days. At the end of the experiment, plants were sampled, weighed and digested with concentrated HNO_3_-H_2_O_2_. We further quantified nitrogen content with the Vario EL elemental analyzer (Elementar Analysensysteme GmbH, Hanau. Germany), while potassium, phosphorus and iron concentrations were measured using inductively coupled plasma atomic emission spectroscopy (710 ICP-OES, Agilent technologies, California, USA) (see [11] for specific details).

***Plant protection experiment***. We set up a separate experiment to examine the effect of bacterial introduction on microbiome-mediated plant protection against plant pathogenic *R. solanacearum* bacterium. To this end, we followed a very similar design as in plant growth-promotion experiment. Briefly, 48 seedling trays were prepared and each inoculated with one *Pseudomonas* consortium. Four additional trays were used as non-inoculated controls. Five days after introduction of *Pseudomonas* consortia, *R. solanacearum* strain QL-Rs1115 [12] was introduced to all 52 trays by soil drenching at a final concentration of 10^6^ CFU g^-1^ soil. Disease severity was assessed individually for each plant every second day after the first plant showed disease symptoms using a disease index scale ranging from 0 to 4 [13], where 0, 1, 2, 3 and 4 refer to no wilt symptoms, 1-25% wilted leaves, 26–50% wilted leaves, 51-75% wilted leaves, and 76-100% of wilted leaves or completely wilted plant, respectively. Disease Severity (DS) was calculated for each tray as DS=[∑(number of diseased plants at this disease level × disease level)/(total number of plants investigated × highest disease level]×100% [14]. See [3] for specific details.

***Quantification of total bacterial and Pseudomonas abundances.*** We used the primer set developed by Almario and colleagues [15], which has specifically been designed to quantitatively amplify the *phlD* gene regardless of the taxonomic identity of the strain. While *phlD* gene could be used for molecular quantification of the total *Pseudomonas* abundances, it could not distinguish the relative abundances of different individual strains [15]. Also, we cannot fully exclude the presence of some PCR bias, it should have not affected our main results because each *Pseudomonas* strain was included equally often within each richness level. As a result, potential strain-specific qPCR biases were the same across richness levels. DNA extracted from the same 0.3 g soil samples used for amplicon sequencing were also used for qPCR quantification of 16S (Eub338: 5'-ACT CCT ACG GGA GGC AGC AG-3' and Eub518: 5'-ACT CCT ACG GGA GGC AGC AG-3' [16] and *PhlD* gene copy numbers (B2BF: 5'-ACC CAC CGC AGC ATC GTT TAT GAG C-3' and B2BR3: 5'-AGC AGA GCG ACG AGA ACT CCA GGG A-3' [15]. qPCR analyses were carried out with Applied Biosystems 7500 Real-Time PCR System (Applied Biosystems, CA, USA) using SYBR Green I fluorescent dye with standard protocol. Briefly, each reaction (20 μL volume) contained 10 μL of SYBR Premix Ex Taq (Takara Biotech. Co., Japan), 2 μL of DNA template and 0.4 μL of both forward and reverse primers (final concentration of 200 nM). The PCR was performed by initially denaturing at 95 °C for 30 s and cycling 40 times with a 5 s denaturisation step at 95 °C. This was followed by a 34 s elongation/extension step at 60 °C and melt curve analysis at 95 °C for 15 s, at 60 °C for 1 min and at 95 °C for 15 s. Each reaction was replicated three times.

***16S rRNA amplicon sequencing:*** The first round of PCR was used to attach the barcodes to primers for MiSeq sequencing, followed by a second round of PCR was then used for paired-end sequencing using Y-shaped adapters. Amplicons were purified using magnetic bead sand, which was denatured with fresh NaOH, beads/PCR reactional volume ratio of 0.8 and final elution volume of 32 μL using Elution Buffer EB (Qiagen). Purified PCR products were quantified using Qubit®3.0 (Life Invitrogen) and amplicon pools were made from twenty-four amplicon samples with different barcodes. The pooled DNA products were used to construct the Illumina Pair-End library following Illumina’s genomic DNA library preparation procedure before paired-end sequencing (2 × 250 base pairs) using Illumina Miseq platform according to the standard protocols.

***DADA2 conformation analysis:*** To ensure the robustness of the OTU bioinformatic pipeline, the 16S rRNA Miseq sequence data was also analyzed using DADA2 pipeline on QIIME2 (version: qiime2-2018.2) where the default overlap length (for merging trimmed paired reads) of forward and reverse reads was decreased to 6 nucleotides [17,18]. Eight nucleotides were removed at the 5’ end of both forward and reverse reads to keep a good sequencing quality at the denoising step. Other parameters were set as trunclen=159, maxEE=1, truncQ=11. With the DADA2 algorithm, taxonomic assignments were resolved with exact sequence features based on amplicon sequence variants (ASVs). Taxonomy was assigned using the Silva database (release 132) at 99% identity [19]. After filtering and rarefaction, on average of 23851 high-quality sequences (min=23,819 and max=23,920) per sample were obtained resulting in a total of 5,522 ASVs, which were included in downstream analysis. MAFFT version 7 (https://mafft.cbrc.jp/alignment/server/phylogeny.html) was used to estimate the phylogeny of all ASVs observed in all samples. We then compared if the richness of inoculated consortium had similar effects on the rare and dominant rhizosphere microbiome taxa depending of the resident microbiome diversity, estimated using both OTUs or ASVs. Qualitatively highly similar results were observed between the two methods (Fig S2 and S3). As a result, the OTU pipeline was used for microbiome diversity analyses, where a slightly lower taxonomic accuracy was traded against a better coverage and representation of rare taxa.

***Calculation of resident microbiome biodiversity****.* Phylogenetic abundance evenness (PAE), an index accounting for both evenness and phylogenetic distribution, which was calculated as follows [20]:

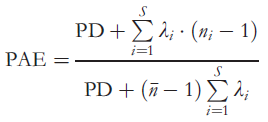

where Faith's Phylogenetic Diversity (PD) is the phylogenetic diversity value which is the sum of all branch lengths represented by the species in the community, λ_i_ is the terminal edge length of OTU i, n_i_ is the abundance of OTU i, S is the number of all OTUs, $\bar{n}$ is the mean of OTU abundance. PD [21] was calculated using the R function pd (include.root=FALSE; package picante) based upon neighbour-joining phylogenetic trees generated with PyNast and FastTree. PD is the sum of the total branch length in a phylogenetic tree for each community and this index contains not only the information of taxonomic diversity but also the phylogenetic relations across whole communities.

**Statistical analyses for supplementary tables and figures**

In order to verify that consortium diversity effects were not driven by the inclusion of particular strains, identity effects were analyzed. To this end, the effect of each of the introduced strain was included as the presence in consortia (binary predictor), and the effect of strain exclusions on consortium richness effects was analyzed [22] (results shown in Tables S5-S7). The effect of consortium diversity and strain identity effects on resident microbiome diversity were tested using four diversity indexes: phylogenetic diversity (PD), OTU richness, Shannon’s diversity and Pielou’s evenness using R package vegan (Table S7).

To assess how ranked OTU abundances correlated with *Pseudomonas* inoculants, we first ranked the abundances of each OTU in the control treatment (without *Pseudomonas* inoculations). We then calculated how OTU abundances correlated with *Pseudomonas* consortium richness using a general linear model, and significantly correlated OTUs were scored as 1, and non-significant as 0. We then used a generalised additive model (GAM) to assess the relationship between ranked OTU abundances and the probability of responding to *Pseudomonas* consortium richness using the R package gam. Separate analyses were carried out for positively and negatively correlated OTUs. We also calculated the correlations between resident bacterial abundances (at phylum or family level) and *Pseudomonas* consortium richness using general linear models (shown in scatter diagrams in Fig. S4A-B). The OTUs whose abundances were significantly increased or decreased with probiotic consortium richness are presented as heatmaps in Fig. S4C-D.

We also used Structural Equation Modelling (SEM) to disentangle whether the effects of probiotic consortia on the plant growth, nutrient assimilation and pathogen tolerance were directly driven by the plant-beneficial functions provided by the inoculated consortia, or indirectly via shifts in the composition and diversity of the resident rhizosphere microbiome (Fig. S5).

***References***

1.	Wei Z, Yang T, Friman V-P, Xu Y, Shen Q, Jousset A. 2015 Trophic network architecture of root-associated bacterial communities determines pathogen invasion and plant health. *Nature Communications* **6**, 8413. (doi:10.1038/ncomms9413)

2.	Ornston LN, Stanier RY. 1966 The Conversion of Catechol and Protocatechuate to β-Ketoadipate by Pseudomonas putida. *Journal of Biological Chemistry* **241**, 3776–3786. (doi:10.1016/S0021-9258(18)99839-X)

3.	Hu J *et al.* 2016 Probiotic diversity enhances rhizosphere microbiome function and plant disease suppression. *mBio* **7**, e01790-16. (doi:10.1128/mBio.01790-16)

4.	Schwyn B, Neilands JB. 1987 Universal chemical assay for the detection and determination of siderophores. *Analytical Biochemistry* **160**, 47–56. (doi:10.1016/0003-2697(87)90612-9)

5.	Ghosh SK, Pal S, Chakraborty N. 2015 The qualitative and quantitative assay of siderophore production by some microorganisms and effect of different media on its production. *International Journal of Chemical Science* **13**, 1621–1629.

6.	Landy M, Warren GH, RosenmanM SB, Colio LG. 1948 Bacillomycin: An Antibiotic from Bacillus subtilis Active against Pathogenic Fungi. *Proceedings of the Society for Experimental Biology and Medicine* **67**, 539–541. (doi:10.3181/00379727-67-16367)

7.	Nautiyal CS. 1999 An efficient microbiological growth medium for screening phosphate solubilizing microorganisms. *FEMS Microbiology Letters* **170**, 265–270. (doi:10.1111/j.1574-6968.1999.tb13383.x)

8.	Tsang S, Phu F, Baum MM, Poskrebyshev GA. 2007 Determination of phosphate/arsenate by a modified molybdenum blue method and reduction of arsenate by S_2_O_4_^2−^. *Talanta* **71**, 1560–1568. (doi:10.1016/j.talanta.2006.07.043)

9.	Elphinstone JG, Hennessy J, Wilson JK, Stead DE. 1996 Sensitivity of different methods for the detection of *Ralstonia solanacearum* in potato tuber extracts. *EPPO Bulletin* **26**, 663–678. (doi:10.1111/j.1365-2338.1996.tb01511.x)

10.	Wei Z, Huang J, Tan S, Mei X, Shen Q, Xu Y. 2013 The congeneric strain *Ralstonia pickettii* QL-A6 of *Ralstonia solanacearum* as an effective biocontrol agent for bacterial wilt of tomato. *Biological Control* **65**, 278–285. (doi:10.1016/j.biocontrol.2012.12.010)

11.	Hu J, Wei Z, Weidner S. 2017 Probiotic *Pseudomonas* communities enhance plant growth and nutrient assimilation via diversity-mediated ecosystem functioning. *Soil Biology and Biochemistry* **113**, 122–129. (doi:10.1016/j.soilbio.2017.05.029)

12.	Hayward AC. 1991 Biology and Epidemiology of Bacterial Wilt Caused by Pseudomonas Solanacearum. *Annual Review of Phytopathology* **29**, 65–87.

13.	Tans-Kersten J, Brown D, Allen C. 2004 Swimming Motility, a Virulence Trait of Ralstonia solanacearum, Is Regulated by FlhDC and the Plant Host Environment. *Molecular Plant-Microbe Interactions* **17**, 686–695. (doi:10.1094/MPMI.2004.17.6.686)

14.	Wei Z, Yang X, Yin S, Shen Q, Ran W, Xu Y. 2011 Efficacy of *Bacillus*-fortified organic fertiliser in controlling bacterial wilt of tomato in the field. *Applied Soil Ecology* **48**, 152–159. (doi:doi.org/10.1016/j.apsoil.2011.03.013)

15.	Almario J, Moënne-Loccoz Y, Muller D. 2013 Monitoring of the relation between 2,4-diacetylphloroglucinol-producing *Pseudomonas* and *Thielaviopsis basicola* populations by real-time PCR in tobacco black root-rot suppressive and conducive soils. *Soil Biology and Biochemistry* **57**, 144–155. (doi:10.1016/j.soilbio.2012.09.003)

16.	Fierer N, Jackson JA, Vilgalys R, Jackson RB. 2005 Assessment of Soil Microbial Community Structure by Use of Taxon-Specific Quantitative PCR Assays. *Applied and Environmental Microbiology* **71**, 4117–4120. (doi:10.1128/AEM.71.7.4117-4120.2005)

17.	Callahan BJ, McMurdie PJ, Rosen MJ, Han AW, Johnson AJA, Holmes SP. 2016 DADA2: High-resolution sample inference from Illumina amplicon data. *Nature Methods* **13**, 581–583. (doi:10.1038/nmeth.3869)

18.	Caporaso JG *et al.* 2010 QIIME allows analysis of high-throughput community sequencing data. *Nature methods* **7**, 335–336. (doi:10.1038/nmeth.f.303)

19.	Quast C, Pruesse E, Yilmaz P, Gerken J, Schweer T, Yarza P, Peplies J, Glöckner FO. 2012 The SILVA ribosomal RNA gene database project: improved data processing and web-based tools. *Nucleic Acids Research* **41**, D590–D596. (doi:10.1093/nar/gks1219)

20.	Cadotte MW, Davies TJ, Regetz J, Kembel SW, Cleland E, Oakley TH. 2010 Phylogenetic diversity metrics for ecological communities: integrating species richness, abundance and evolutionary history. *Ecology Letters* **13**, 96–105. (doi:10.1111/j.1461-0248.2009.01405.x)

21.	Faith DP. 1992 Conservation evaluation and phylogenetic diversity. *Biological Conservation* **61**, 1–10. (doi:10.1016/0006-3207(92)91201-3)

22.	Jousset A, Schulz W, Scheu S, Eisenhauer N. 2011 Intraspecific genotypic richness and relatedness predict the invasibility of microbial communities. *The ISME Journal* **5**, 1108–1114. (doi:10.1038/ismej.2011.9)
